# Presynaptic NMDA receptors facilitate short-term plasticity and BDNF release at hippocampal mossy fiber synapses

**DOI:** 10.1101/2021.01.21.427714

**Authors:** Pablo J. Lituma, Hyung-Bae Kwon, Karina Alviña, Rafael Lujan, Pablo E. Castillo

## Abstract

Neurotransmitter release is a highly controlled process by which synapses can critically regulate information transfer within neural circuits. While presynaptic receptors –typically activated by neurotransmitters and modulated by neuromodulators– provide a powerful way of fine-tuning synaptic function, their contribution to activity-dependent changes in transmitter release remains poorly understood. Here, we report that presynaptic NMDA receptors (preNMDARs) at mossy fiber boutons in the rodent hippocampus can be activated by physiologically relevant patterns of activity and selectively enhance short-term synaptic plasticity at mossy fiber inputs onto CA3 pyramidal cells and mossy cells, but not onto inhibitory interneurons. Moreover, preNMDARs facilitate brain-derived neurotrophic factor (BDNF) release and contribute to presynaptic calcium rise. Taken together, our results indicate that by increasing presynaptic calcium, preNMDARs fine tune mossy fiber neurotransmission and can control information transfer during dentate granule cell burst activity that normally occur *in vivo*.

## Introduction

Neurotransmission is a dynamic and highly regulated process. The activation of ionotropic and metabotropic presynaptic autoreceptors provides a powerful way of fine-tuning neurotransmission via the facilitation or inhibition of neurotransmitter release (Burke & Bender, 2019; Engelman & MacDermott, 2004; Miller, 1998; Pinheiro & Mulle, 2008; Schicker et al., 2008). Due to their unique functional properties, including high calcium-permeability, slow kinetics and well-characterized role as coincidence-detectors (Cull-Candy et al., 2001; Lau & Zukin, 2007; Paoletti et al., 2013; Traynelis et al., 2010), presynaptic NMDA receptors (preNMDARs) have received particular attention (Banerjee et al., 2016; Bouvier et al., 2015; Bouvier et al., 2018; Duguid, 2013; Duguid & Smart, 2009; Wong et al., 2020). Regulation of neurotransmitter release by NMDA autoreceptors in the brain was suggested three decades ago (Martin et al., 1991). Anatomical evidence for preNMDARs arose from an immuno-electron microscopy study revealing NMDARs at the mossy fiber giant bouton of the monkey hippocampus (Siegel et al., 1994), followed by functional studies in the entorhinal cortex indicating that preNMDARs tonically increase spontaneous glutamate release and also facilitate evoked release in a frequency-dependent manner (Berretta & Jones, 1996; Woodhall et al., 2001). Since these early studies, although evidence for preNMDARs has accumulated throughout the brain (Banerjee et al., 2016; Bouvier et al., 2018; Duguid & Smart, 2009), the presence and functional relevance of preNMDARs at key synapses in the brain have been called into question (Carter & Jahr, 2016; Duguid, 2013).

Mossy fibers (mf) – the axons of dentate granule cells (GCs) – establish excitatory synapses onto proximal dendrites of CA3 pyramidal neurons, thereby conveying a major excitatory input to the hippocampus proper (Amaral et al., 2007; Henze et al., 2000). This synapse displays uniquely robust frequency facilitation both *in vitro* (Nicoll & Schmitz, 2005; Salin et al., 1996; Vyleta et al., 2016) and *in vivo* (Hagena & Manahan-Vaughan, 2010; Vandael et al., 2020). The molecular basis of this short-term plasticity is not fully understood but likely relies on diverse presynaptic mechanisms that increase glutamate release (Jackman & Regehr, 2017; Rebola et al., 2017). Short-term, use-dependent facilitation is believed to play a critical role in information transfer, circuit dynamics and short-term memory (Abbott & Regehr, 2004; Jackman & Regehr, 2017; Klug et al., 2012). The mf-CA3 synapse can strongly drive the CA3 network during short bursts of presynaptic activity (Chamberland et al., 2018; Henze et al., 2002; Vyleta et al., 2016; Zucca et al., 2017), an effect that likely results from two key properties of this synapse, namely, its strong frequency facilitation and proximal dendritic localization. In addition to CA3 pyramidal neurons, mf axons establish synaptic connections with hilar mossy cells (MC) and inhibitory interneurons (IN) (Amaral et al., 2007; Henze et al., 2000). These connections also display robust short-term plasticity (Lysetskiy et al., 2005; Toth et al., 2000), which may contribute significantly to information transfer and dynamic modulation of the dentate gyrus (DG)-CA3 circuit (Bischofberger et al., 2006; Evstratova & Toth, 2014; Lawrence & McBain, 2003). Despite early evidence for preNMDARs at mf boutons (Siegel et al., 1994), whether these receptors modulate neurotransmission at mf synapses is unknown. Intriguingly, mfs contain one of the highest expression levels of brain-derived neurotrophic factor, BDNF (Conner et al., 1997). While preNMDARs were implicated in BDNF release at corticostriatal synapses (Park et al., 2014), whether putative preNMDARs impact BDNF release at mf synapses remains unexplored.

Here, to examine the potential presence and impact of preNMDARs at mf synapses, we utilized multiple approaches, including immunoelectron microscopy, selective pharmacology for NMDARs, a genetic knockout strategy to remove NMDARs from presynaptic GCs, two-photon imaging of BDNF release, and presynaptic Ca^2+^ signals in acute rodent hippocampal slices. Our findings indicate that preNMDARs contribute to mf short-term plasticity and promotes BDNF release likely by increasing presynaptic Ca^2+^. Thus, preNMDARs at mfs may facilitate information transfer and provide an important point of regulation in the DG – CA3 circuit by regulating both glutamate and BDNF release.

## Results

### Electron microscopy reveals presynaptic NMDA receptors at mossy fiber terminals

To determine the potential localization of NMDA receptors at the mf terminals of the rodent hippocampus, we performed electron microscopy and post-embedding immunogold labeling in rats using a validated antibody for the obligatory subunit GluN1 (Petralia et al., 1994; Siegel et al., 1994; Takumi et al., 1999; Watanabe et al., 1998). Gold particles were detected in the main body of the postsynaptic density as well as presynaptic mf terminals (**Figure 1A-C**). GluN1 localized in mf boutons in a relatively high proportion to the active zone, as compared to associational-commissural (ac) synapse in the same CA3 pyramidal neuron (**Figure 1D**; mf, ∼32% presynaptic particles; ac, <10% presynaptic particles; n = 3 animals). Similar quantification for AMPA receptors did not reveal presynaptic localization of these receptors in either mf or associational commissural synapses (**Figure 1-figure supplement 1**; ∼5% presynaptic particles, n = 3 animals). Together, these results provide anatomical evidence for preNMDARs at mf-CA3 synapses.

**Figure 1.**
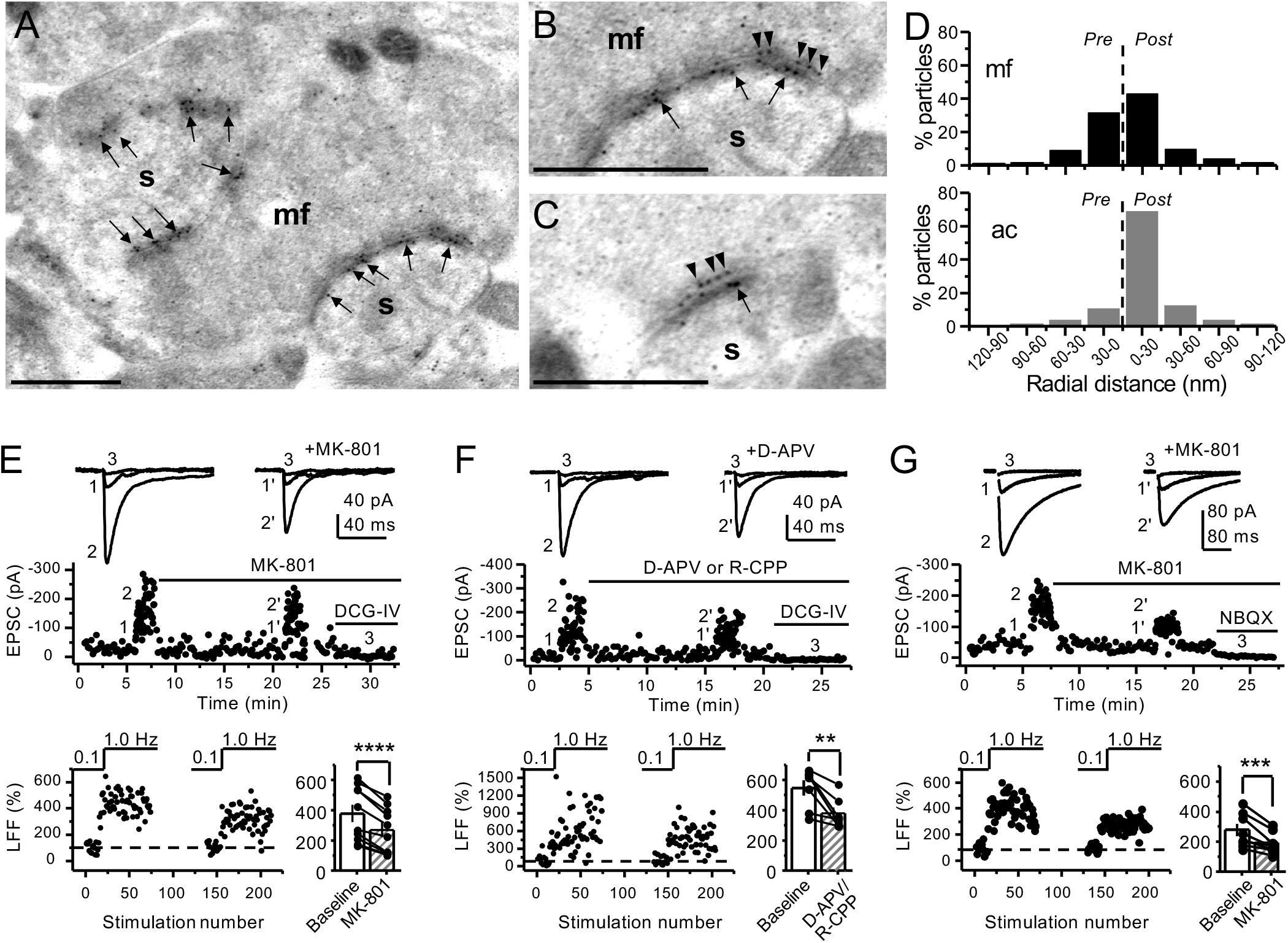
Anatomical and functional evidence for preNMDARs at mossy fiber synapses. (**A**) Image of a mossy fiber (mf) giant bouton and postsynaptic spines (s). (**B, C**) Higher magnification of mf synapses. Arrows indicate postsynaptic GluN1 whereas arrowheads indicate presynaptic GluN1. Calibration bars: 500 nm. (**D**) Mossy fiber (mf) and associational commissural (ac) synaptic GluN1 immuno-particle radial distribution (30 nm bins), mf: 34 synapses, 100 presynaptic particles; ac: 25 synapses, 24 presynaptic particles; 3 animals. (**E**) AMPAR-ESPCs were recorded at V_h_= -70 mV in the presence of 0.5 µM LY303070 and 100 µM picrotoxin. Low-frequency facilitation (LFF), induced by stepping stimulation frequency from 0.1 to 1 Hz, was assessed before and after bath application of MK-801 (50 µM). MK-801 significantly reduced LFF (baseline 378 ± 57%, MK-801 270 ± 48%, n = 10 cells, 9 animals; baseline vs MK-801, p = 3.8 x 10^-5^, paired *t-*test). In all panels of this figure: representative traces (*top*), representative experiment (*middle*), normalized LFF and summary plot (*bottom*). DCG-IV (1 µM) was applied at the end of all recordings to confirm mf-CA3 transmission. (**F**) D- APV (100 µM) or R-CPP (50 µM) application also reduced LFF (baseline 546 ± 50%, D-APV/R- CPP 380 ± 38%, n = 7 cells, 5 animals; baseline vs D-APV/R-CPP, p = 0.00743, paired *t-*test). (**G**) KAR-EPSCs were recorded at V_h_= -70 mV in the presence of 15 µM LY303070 and 100 µM picrotoxin. In addition, NMDAR-mediated transmission was blocked intracellularly by loading MK-801 (2 mM) in the patch-pipette. Bath application of MK-801 (50 µM) significantly reduced LFF (baseline 278 ± 40%, MK-801 195 ± 26% n = 8 cells, 6 animals; baseline vs MK-801, p = 0.00259, paired *t*-test). Data are presented as mean ± s.e.m. ** p < 0.01; *** p < 0.005; **** p < 0.001.

### Both NMDAR antagonism and genetic deletion from presynaptic granule cells reduce mossy fiber low-frequency facilitation

Presynaptic short-term plasticity, in the form of low-frequency (∼1 Hz) facilitation (LFF), is uniquely robust at the mf-CA3 synapse (Nicoll & Schmitz, 2005; Salin et al., 1996). To test a potential involvement of preNMDARs in LFF, we monitored AMPAR-mediated excitatory postsynaptic currents (EPSCs) from CA3 pyramidal neurons in acute rat hippocampal slices.

Neurons were held at V_h_= -70 mV to minimize postsynaptic NMDAR conductance, and mfs were focally stimulated with a bipolar electrode (theta glass pipette) placed in *stratum lucidum* ∼100 µm from the recorded cell. LFF was induced by stepping the stimulation frequency from 0.1 Hz to 1 Hz for ∼2 min in the presence of picrotoxin (100 µM) to block fast inhibitory synaptic transmission, and a low concentration of the AMPAR noncompetitive antagonist LY303070 (0.5 μM) to minimize CA3-CA3 recurrent activity (Kwon & Castillo, 2008). Bath-application of the NMDAR irreversible open channel blocker MK-801 (50 μM) significantly reduced LFF (**Figure 1E**). In addition, the competitive NMDAR antagonists D-APV (100 µM) or R-CPP (50 µM) yielded a comparable reduction of facilitation (**Figure 1F**). To confirm that these synaptic responses were mediated by mfs, the mGluR2/3 agonist DCG-IV (1 µM) was applied at the end of all recordings (Kamiya et al., 1996). To control for stability, we performed interleaved experiments in the absence of NMDAR antagonists and found that LFF remained unchanged (**Figure 1-figure supplement 2A**). These findings indicate NMDAR antagonism reduces mf- CA3 short-term plasticity (LFF), suggesting that preNMDARs could contribute to this form of presynaptic plasticity.

The reduction in facilitation of AMPAR-transmission could be due to dampening of CA3 recurrent activity by NMDAR antagonism (Henze et al., 2000; Kwon & Castillo, 2008; Nicoll & Schmitz, 2005). To discard this possibility, we repeated our experiments in a much less excitable network in which AMPAR-mediated synaptic transmission was selectively blocked by a high concentration of the noncompetitive antagonist LY303070 (15 μM) and monitored the kainate receptor (KAR)-mediated component of mf synaptic transmission (Castillo et al., 1997; Kwon & Castillo, 2008). In addition, 2 mM MK-801 was included in the intracellular recording solution to block postsynaptic NMDARs (Corlew et al., 2008) (**Figure 1-figure supplement 3**). To further ensure postsynaptic NMDAR blockade, we voltage-clamped the CA3 pyramidal neuron at -70 mV and waited until NMDAR-mediated transmission was eliminated and only KAR-EPSCs remained. Under these recording conditions, bath-application of MK-801 (50 μM) also reduced LFF of KAR-mediated transmission (**Figure 1G**), whereas LFF remained unchanged in interleaved control experiments (**Figure 1-figure supplement 2B**). At the end of these recordings, 10 μM NBQX was applied to confirm KAR-transmission (**Figure 1G; Figure 1-figure supplement 2B**) (Castillo et al., 1997; Kwon & Castillo, 2008). It is therefore unlikely that the reduction of LFF mediated by NMDAR antagonism could be explained by recurrent network activity, suggesting a direct effect on transmitter release.

To further support a role of preNMDARs in mf LFF, we took a genetic approach by conditionally removing NMDARs from GCs in *Grin1* floxed mice. To this end, an AAV5-CamKII-Cre-GFP virus was bilaterally injected in the DG to selectively delete *Grin1* expression, whereas AAV5-CamKII-eGFP was injected in littermates as a control at postnatal days 16-20 in both groups (**Figure 2A**). Two weeks after surgery, we prepared acute hippocampal slices and examined the efficacy of *Grin1* deletion by analyzing NMDAR-mediated transmission in GFP^+^ GCs of *Grin1*-cKO and control mice. We confirmed that in contrast to control mice, no NMDAR-EPSCs were elicited by electrically stimulating medial perforant path inputs in *Grin1*-cKO GCs voltage-clamped at +40 mV in the presence of 100 μM picrotoxin and 10 μM NBQX (**Figure 2B**). As expected, the NMDAR/AMPAR ratio was significantly reduced in *Grin1*-cKO mice compared to control (**Figure 2C**). Only acute slices that exhibited robust GFP fluorescence in the DG were tested for LFF of AMPAR-transmission in CA3. We found that LFF was significantly reduced in *Grin1*-cKOs as compared to controls (**Figure 2D**), indicating that genetic removal of NMDARs from GCs recapitulated NMDAR antagonism (**Figure 1E-G**). *Grin1* deletion did not affect basal transmitter release as indicated by a comparable paired-pulse ratio to control (Control: 2.5 ± 0.36, n = 13 cells; *Grin1* cKO: 2.4 ± 0.31, n = 13 cells; U > 0.5, Mann-Whitney test). Collectively, our findings using two distinct approaches strongly suggest that NMDAR activation in GCs increases LFF of mf-CA3 synaptic transmission.

**Figure 2.**
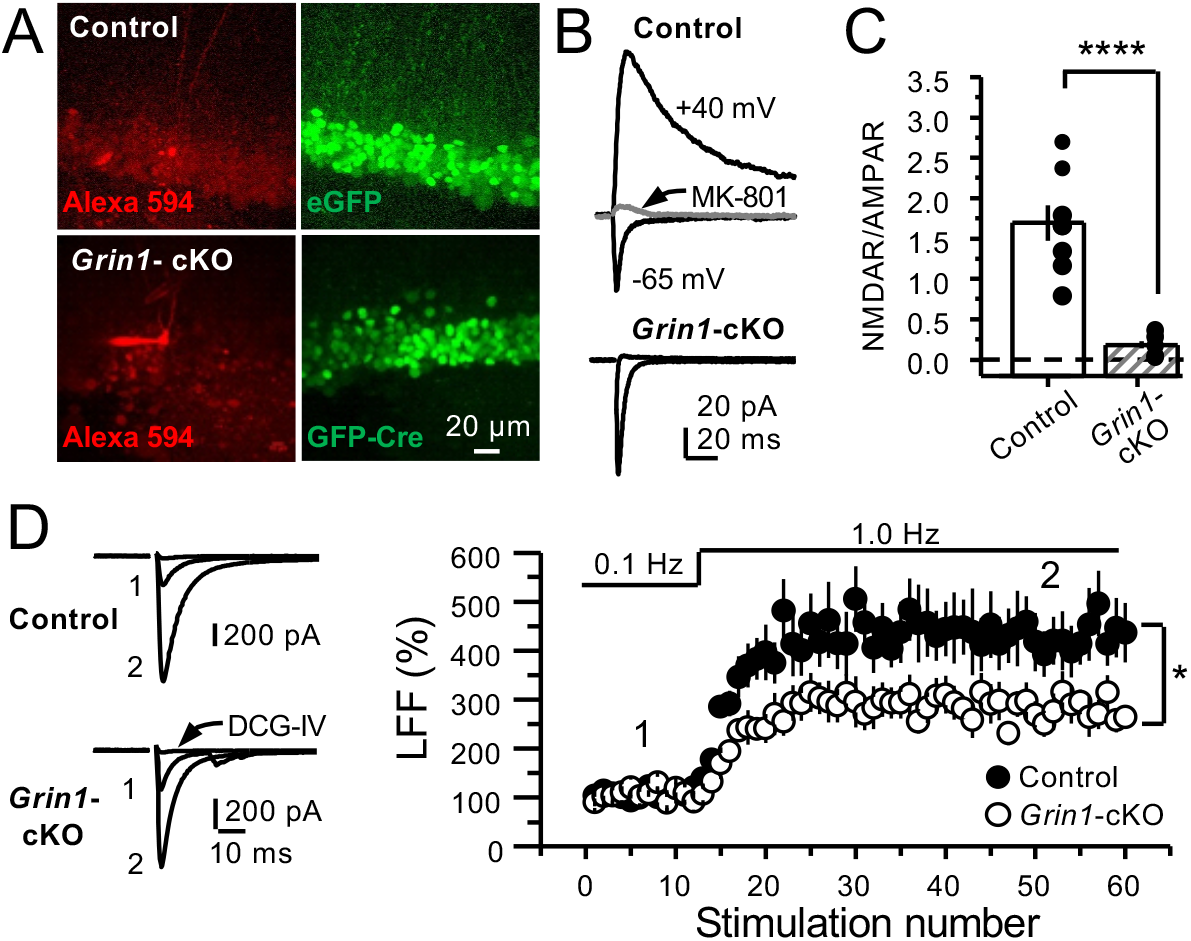
GluN1 deletion from granule cells reduces mf-CA3 facilitation. (**A**) Representative images showing GCs patch-loaded with Alexa 594 (35 µM) *(left),* and GFP expression in GCs *(right)*. (**B**) Representative EPSCs recorded from control (GFP^+^) and *Grin1*- cKO (Cre-GFP^+^) GCs. Synaptic responses were elicited by activating medial perforant-path inputs. AMPAR-ESPCs were recorded at V_h_= -65 mV in the presence of 100 µM picrotoxin, NMDAR-EPSCs were isolated with 10 µM NBQX and recorded at +40 mV. MK-801 (20 µM) was applied at the end of each recording. (**C**) Summary plot demonstrating that GluN1 deletion from GCs virtually abolished NMDAR-mediated transmission indicated by a strong reduction of NMDAR/AMPAR in *Grin1*-cKO GCs as compared to controls (control 1.61 ± 0.18, n = 9 cells, 9 animals, *Grin1*-cKO 0.18 ± 0.04, n = 10 cells, 10 animals; control vs *Grin1*-cKO, p = 9.2 x 10^-6^, unpaired *t*-test). (**D**) LFF was significantly reduced in GluN1-deficient animals (control, 430 ± 5 %, n = 13 cells, 10 animals; *Grin1*-cKO, 291 ± 6 %, n = 11 cells, 10 animals; p = 0.0239, unpaired *t*-test). Representative traces (*left*) and summary plot (*right*). LFF was induced by stepping stimulation frequency from 0.1 to 1 Hz. DCG-IV (1 µM) was added at the end of each experiment. Data are presented as mean ± s.e.m. * p < 0.05; **** p < 0.001.

### Reduced facilitation by NMDAR antagonism is independent of the granule cell somatodendritic compartment

Bath application of MK-801 could have blocked dendritic NMDARs in GCs and potentially affected transmitter release (Christie & Jahr, 2008; Duguid, 2013). To address this possibility, we repeated our experiments after performing a surgical cut in the granular layer of the DG in order to isolate mf axons from GCs (**Figure 3-figure supplement 1A**). Under these conditions, MK-801 bath application still reduced LFF (**Figure 3A**), and LFF was stable in control, acutely transected axons (**Figure 3B**). In addition, puffing D-APV (2 mM) in *stratum lucidum* near (∼200 µm) the recorded neuron also reduced LFF (**Figure 3C**), whereas puffing ACSF had no effect (**Figure 3D**). Lastly, in a set of control experiments, we confirmed that D-APV puffs were sufficient to transiently block NMDAR-mediated transmission in CA3 but not in DG (**Figure 3-figure supplement 1B,C**). Together, these results support the notion that LFF reduction was due to the blockade of preNMDARs but not somatodendritic NMDARs on GCs.

**Figure 3.**
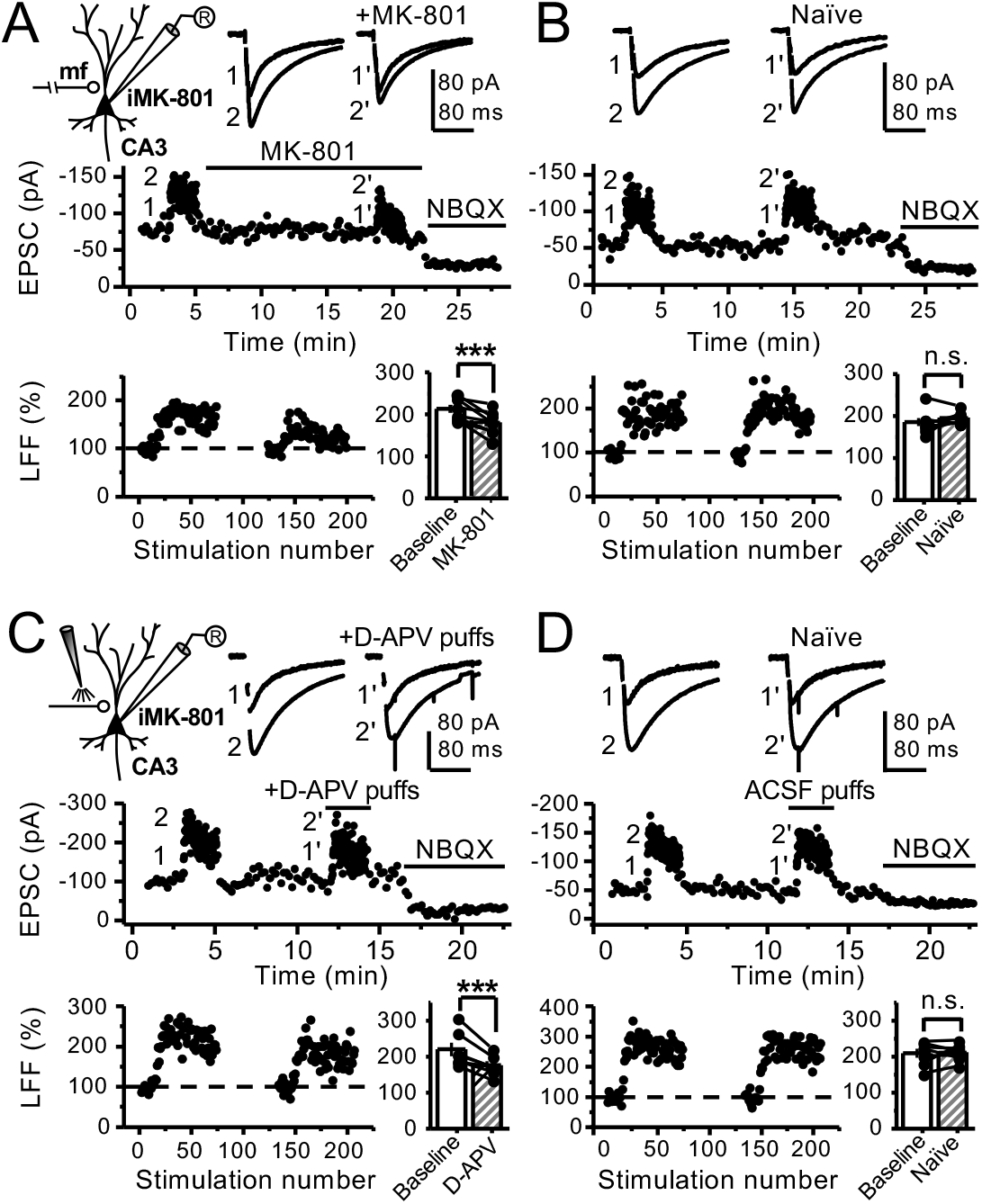
Reduced facilitation by NMDAR antagonism is independent of the GC somatodendritic compartment. (**A**) KAR-EPSCs were recorded at V_h_= -70 mV in the presence of 15 µM LY303070 and 100 µM picrotoxin. In addition, NMDAR-mediated transmission was blocked intracellularly by loading MK-801 (2 mM) in the patch-pipette. LFF of KAR-EPSCs was assessed as in Fig. 1G but with transected mf axons (see Methods). Bath application of MK-801 (50 µM) significantly reduced LFF (baseline 213 ± 9%, MK-801 181 ± 10%, n = 8 cells, 7 animals; baseline vs MK-801, p = 0.002, paired *t*-test). In all panels of this figure: recording arrangement (*inset*), representative traces (*top*), representative experiment (*middle*), normalized LFF and summary plot (*bottom*). (**B**) Stable LFF in transected, naïve slices (baseline 186 ± 10%, naïve 196 ± 5%, n = 8 cells, 7 animals; baseline vs naïve, p = 0.278, paired *t*-test). (**C**) LFF was induced before and during puff application of D-APV (2 mM) in *stratum lucidum*. This manipulation significantly reduced facilitation (baseline 220 ± 19%, D-APV puff 176 ± 11%, n = 7 cells, 7 animals; baseline vs D-APV puff, p = 0.003, paired *t*-test). (**D**) Stable LFF in acute slices during puff application of ACSF (baseline 210% ± 12, naïve 213% ± 9, n = 7 cells, 7 animals; baseline vs naïve, p = 0.778, paired *t*-test). NBQX (10 µM) was applied at the end of all recordings to confirm mf KAR transmission. Data are presented as mean ± s.e.m. *** p < 0.005.

### PreNMDARs boost information transfer by enhancing burst-induced facilitation at mossy fiber synapses

GCs *in vivo* typically fire in brief bursts (Diamantaki et al., 2016; GoodSmith et al., 2017; Henze et al., 2002; Pernia-Andrade & Jonas, 2014; Senzai & Buzsaki, 2017). To test whether preNMDARs contribute to synaptic facilitation that occurs during more physiological patterns of activity, mfs were activated with brief bursts (5 stimuli, 25 Hz). We first took an optogenetic approach and used a Cre-dependent ChIEF virus to selectively light-activate mf-CA3 synapses in *Grin1*-cKO and control mice. Thus, animals were injected with a mix of AAV5-CamKII-CreGFP + AAV-DJ-FLEX-ChIEF-tdTomato viruses in the DG (**Figure 4A**). At least four weeks after surgery, acute slices were prepared and burst-induced facilitation of AMPAR-mediated transmission in CA3 was assessed (**Figure 4B,C**). Burst-induced facilitation triggered by light stimulation and measured as the ratio of EPSCs elicited by the 5^th^ and 1^st^ pulse (P5/P1 ratio), was significantly reduced in *Grin1*-cKO animals as compared to controls. Because these bursts of activity can activate the CA3 network (Henze et al., 2000; Kwon & Castillo, 2008; Nicoll & Schmitz, 2005), we next monitored KAR-EPSCs under conditions of low excitability (as in Figure 1G). MK-801 bath application also reduced burst-induced facilitation, whereas facilitation remained unchanged in naïve slices (**Figure 4D,E**). In a separate set of experiments, we confirmed the reduction of MK-801 on burst-induced facilitation under more physiological recording conditions (**Figure 4-figure supplement 1**). Lastly, we tested whether preNMDARs, by facilitating glutamate release during bursting activity, could bring CA3 pyramidal neurons to threshold and trigger postsynaptic action potentials. To test this possibility, we monitored action potentials elicited by KAR-EPSPs (resting membrane potential -70 ± 2 mV) from CA3 pyramidal neurons intracellularly loaded with 2 mM MK-801. Under these recording conditions, MK-801 bath application significantly reduced the mean number of spikes per burst (**Figure 4F**). No changes in mean spikes per burst were observed in naïve slices over time (**Figure 4G**). Application of 10 μM NBQX at the end of these experiments confirmed that action potentials were induced by KAR-mediated synaptic responses. Consistent with these observations, MK- 801 also reduced the mean number of spikes per burst when AMPAR-mediated action potentials were recorded from CA3 pyramidal neurons (**Figure 4-figure supplement 2**). In control experiments, we found that intracellular MK-801 effectively blocked postsynaptic NMDAR transmission during burst stimulation (**Figure 4-figure supplement 3**). Altogether, these results indicate that preNMDARs at mf-CA3 synapses can contribute to information transfer from the DG to CA3.

**Figure 4.**
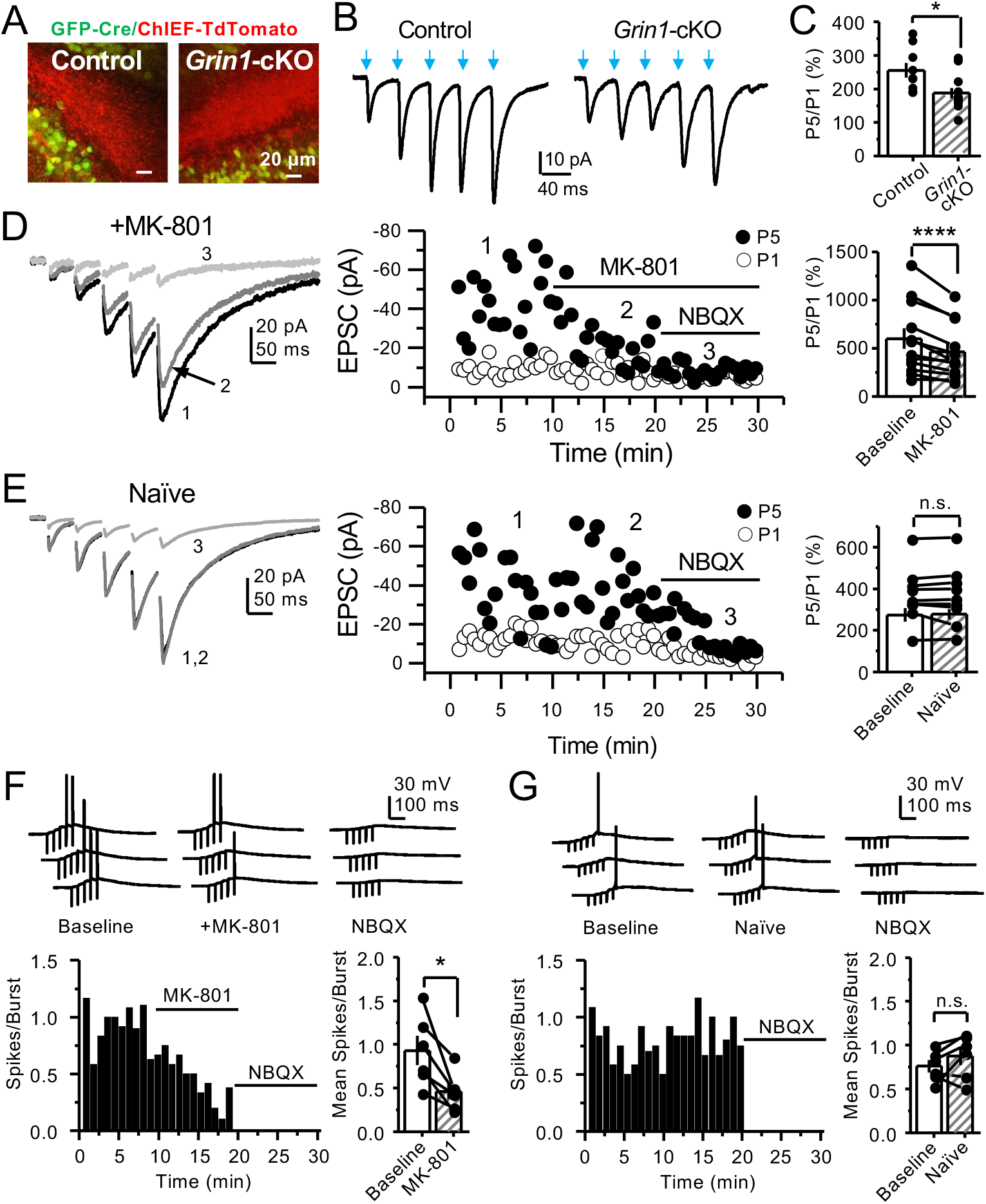
PreNMDARs contribute significantly to burst-induced facilitation and spike transfer. (**A**) Representative images showing expression of GFP-Cre *(left)* and ChIEF- tdTomato *(right)* in the DG of control and *Grin1*-cKO animals. (**B**) Representative AMPAR- EPSCs from control *(left)* and *Grin1*-cKO *(right)* CA3 pyramidal neurons recorded at V_h_= -65 mV and evoked by optical burst-stimulation (5 pulses at 25 Hz) of *stratum lucidum*. Blue arrows indicate light stimulation. (**C**) Summary plot of burst-induced facilitation measured as P5/P1 ratio of optical responses; facilitation was significantly reduced in *Grin1*-cKO animals as compared to control (*Grin1*-cKO 187 ± 16%, n = 12 cells, 9 animals; control 255 ± 22%, n = 9 cells, 8 animals; *Grin1*-cKO vs control, p = 0.0167, unpaired *t*-test). (**D**) Burst-stimulation induced KAR-EPSCs were isolated and recorded as described in Fig. 3, bath-application of MK- 801 (50 µM) significantly reduced facilitation (baseline 601 ± 107%, MK-801 464 ± 84%, n = 13 cells, 10 animals; baseline vs MK-801, p = 0.00042, paired *t*-test). In panels D and E of this figure: representative traces (*left*), representative experiment (*middle*), and summary plot (*right*). (**E**) Burst-induced facilitation was stable in interleaved, naïve slices (baseline 369 ± 45%, naïve 367 ± 48%, n = 9 cells, 9 animals; p = 0.863, paired *t*-test). (**F**) Bath-application of MK-801 (50 µM) reduced KAR-mediated action potentials induced by burst-stimulation (baseline 0.93 ± 0.17, MK-801 0.46 ± 0.09, n = 6 cells, 5 animals; p = 0.036, Wilcoxon Signed Ranks test). In panels F and G of this figure: representative traces (*top)*, representative experiment and summary plot (*bottom)*. (**G**) Stable KAR-mediated action potentials in interleaved naïve slices (baseline 0.76 ± 0.07, naïve 0.88 ± 0.1, n = 6 cells, 5 animals; p = 0.2084, Wilcoxon Signed Ranks test). NBQX (10 µM) was applied at the end of all experiments in panels D-G. Data are presented as mean ± s.e.m. * p < 0.05; **** p < 0.001.

### PreNMDARs contribute to presynaptic calcium rise and can be activated by glutamate

PreNMDARs could facilitate glutamate and BDNF release by increasing presynaptic Ca^2+^ rise (Bouvier et al., 2016; Buchanan et al., 2012; Corlew et al., 2008; Park et al., 2014). To test this possibility at mf-CA3 synapses, we combined a conditional knockout strategy with Ca^2+^ imaging using two-photon laser scanning microscopy. We first deleted preNMDARs by injecting AAV5- CamKII-mCherry-Cre virus in the DG of *Grin1* floxed mice, and littermate animals injected with AAV5-CamKII-mCherry virus served as control (**Figure 5A**). Two weeks after surgery, we confirmed the efficacy of *Grin1* deletion by activating medial perforant path inputs and monitoring NMDAR/AMPAR ratios in GCs of control and *Grin1*-cKO animals (**Figure 5A**).

**Figure 5.**
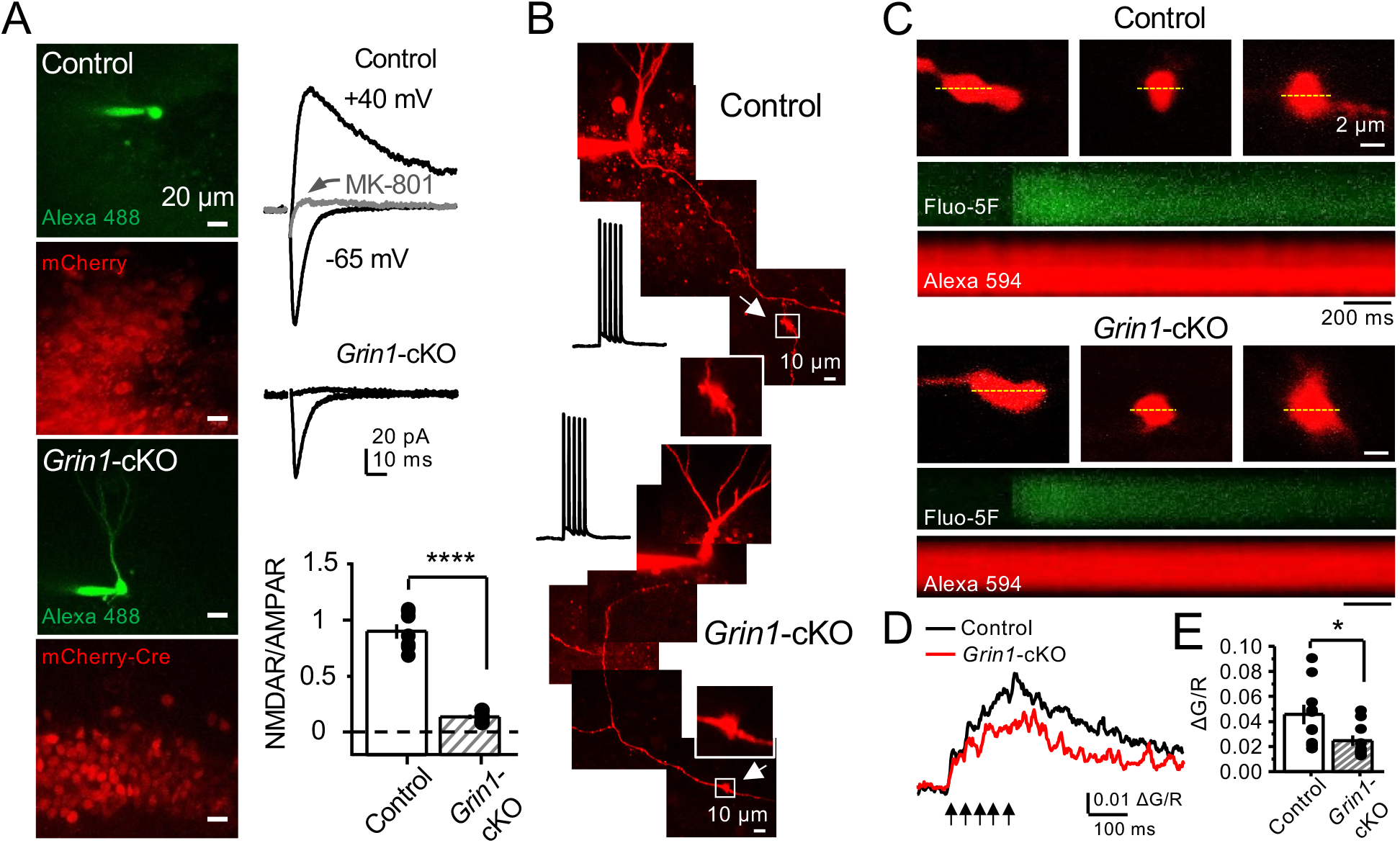
preNMDARs contribute to presynaptic Ca^2+^ rise. (**A**) Representative images showing GCs patch-loaded with Alexa 488 (35 µM) to confirm expression of mCherry *(bottom)*. Representative AMPAR-EPSCs recorded from control *(top)* or *Grin1*-cKO *(middle)* GCs. Synaptic responses were elicited by activating medial perforant-path inputs. AMPAR-ESPCs were recorded at V_h_= -65 mV in the presence of 100 µM picrotoxin, NMDAR-EPSCs were isolated with 10 µM NBQX and recorded at +40 mV. MK-801 (20 µM) was applied at the end of each experiment. Summary plot *(bottom)* demonstrating that GluN1 deletion from GCs virtually abolished NMDAR-mediated transmission indicated by a strong reduction of NMDAR/AMPAR in *Grin1*-cKO granule cells as compared to controls (control 0.90 ± 0.17, n = 7 cells, 6 animals; *Grin1*-cKO 0.13 ± 0.05, n = 6 cells, 6 animals; control vs *Grin1*-cKO, p = 3.81 x 10^-7^, unpaired *t*- test). (**B**) Representative control and *Grin1*-cKO GCs patch-loaded with Fluo-5F (200 µM) and Alexa 594 (35 µM). Arrows indicate the identification of a mf giant bouton, magnified images in white box. (**C**) Three representative mf boutons *(top)* and line scan analysis of calcium transients (CaTs) elicited by 5 action potentials at 25 Hz *(middle,* Fluo-5F) and morphological dye (*bottom*, Alexa 594), in Control and *Grin1*-cKO animals. Dotted line (yellow) indicates line scan location. Red Channel, Alexa 594; Green Channel, Fluo-5F. (**D, E**) Peak analysis of the 5^th^ pulse ΔG/R revealed a significant reduction in Ca^2+^ rise of *Grin1*-cKO animals as compared to Control (control 0.046 ± 0.01, n = 10 boutons, 8 animals; *Grin1*-cKO 0.025 ± 0.004, n = 10 boutons, 8 animals; control vs. *Grin1*-cKO, U = 0.017, Mann-Whitney test). Arrows indicate mf activation. Data are presented as mean ± s.e.m. * U < 0.05; **** p < 0.001.

Virtually no NMDAR-EPSCs were detected at V_h_= +40 mV in *Grin1*-cKO animals (**Figure 5A**). Acute slices that exhibited robust mCherry fluorescence in the DG were used for Ca^2+^ imaging experiments. To maximize our ability to detect preNMDAR-mediated Ca^2+^ signals, we used a recording solution that contained 0 mM Mg^2+^, 4 mM Ca^2+^ and 10 μM D-Serine (Carter & Jahr, 2016). GCs expressing mCherry were patch-loaded with 35 µM Alexa 594 (used as morphological dye) and 200 µM Fluo-5F, and mf axons were imaged and followed towards CA3 until giant boutons (white arrows) were identified (**Figure 5B**). We found that Ca^2+^ transients (CaTs) elicited by direct current injection in the GC soma (5 action potentials, 25 Hz) were significantly smaller in *Grin1*-cKO animals as compared to control (**Figure 5C-E**). In addition, NMDAR antagonism with D-APV reduced presynaptic Ca^2+^ rise even under more physiological Mg^+2^ concentration in acute rat hippocampal slices (**Figure 5-figure supplement 1**). Thus, preNMDARs contribute significantly to presynaptic Ca^2+^ rise in mf boutons, and by this means likely facilitates synaptic transmission, although a potential contribution of Ca^2+^ rise-independent effects cannot be discarded.

Lastly, we sought to determine if direct activation of preNMDARs could drive Ca^2+^ influx in mf giant boutons. To test this possibility, we elicited CaTs by two-photon glutamate uncaging (2PU) on mf boutons of control and *Grin1*-cKO animals (**Figure 6A**). As previously described, mCherry GCs were patch-loaded with Alexa 594 and Fluo-5F in a recording solution designed to maximize the detection of preNMDAR-mediated Ca^2+^ signals (as in Figure 5). We first confirmed that glutamate 2PU-induced CaTs in dendritic spine heads of GCs were strongly reduced in *Grin1*-cKO animals as compared to controls (**Figure 6B,C**). To verify that reduced Ca^2+^ signals (ΔG/R) were a result of *Grin1* deletion and not differences in uncaging laser power, we performed a laser power intensity–response curve, and found that *Grin1*-cKO animals exhibited reduced ΔG/R signals as compared to control regardless of laser power intensity (**Figure 6-figure supplement 1**). We next measured glutamate 2PU-induced CaTs in mf giant boutons (identified as in Figure 5B) and found that single uncaging pulses were insufficient to drive detectable CaTs in control boutons (**Figure 6-figure supplement 2**). However, a burst of 2PU stimulation (5 pulses, 25 Hz) induced CaTs in mf boutons of control but not in *Grin1*-cKO animals (**Figure 6D,E**). Additionally, CaTs elicited by 2PU stimulation were abolished by D-APV application (**Figure 6-figure supplement 3**). These findings indicate that brief bursts of glutamate 2PU, a manipulation that mimics endogenous release of glutamate during physiological patterns of activity, induces presynaptic Ca^2+^ influx in mf boutons by activating preNMDARs.

**Figure 6.**
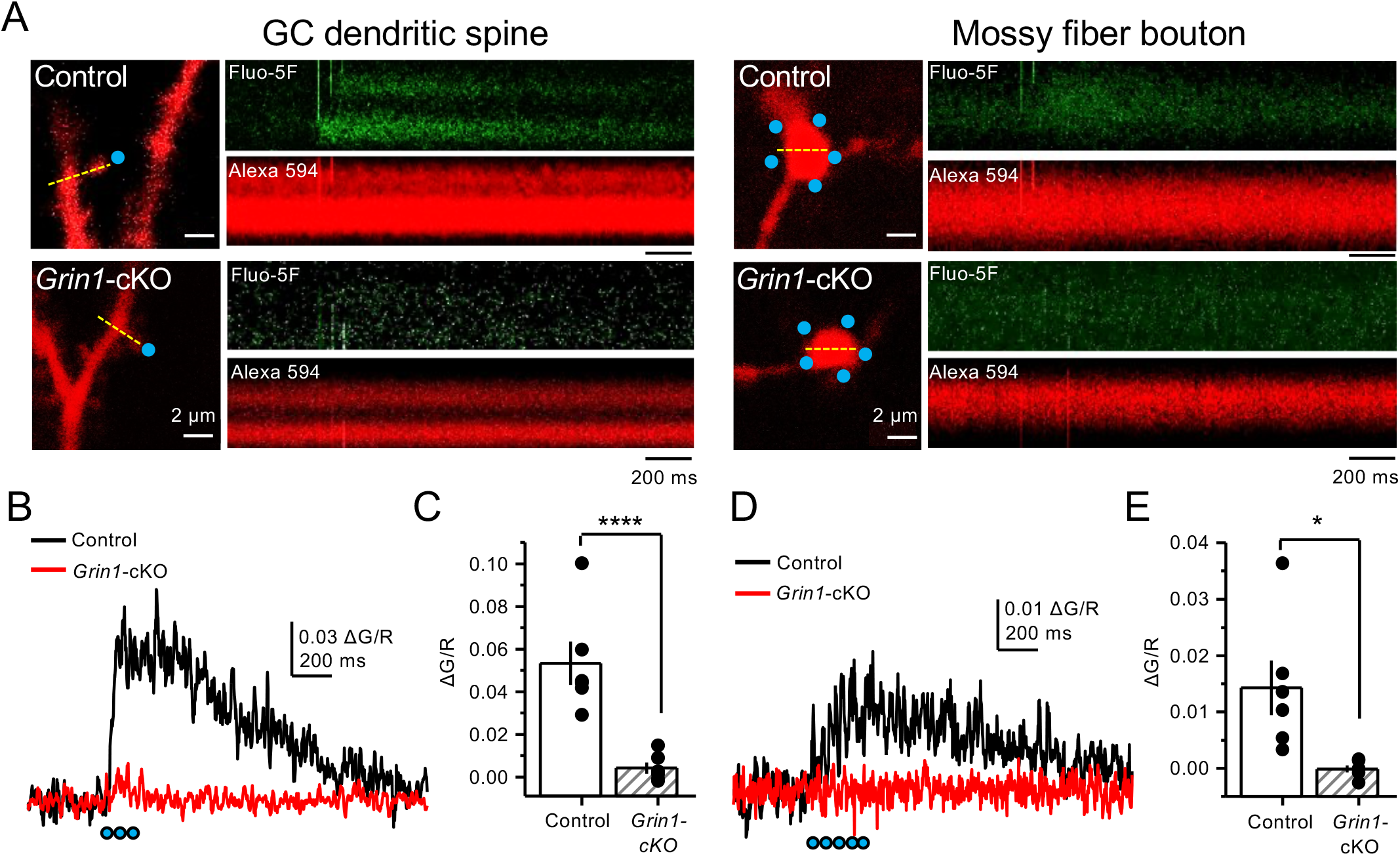
Uncaging glutamate induces Ca^2+^ rise mossy fiber boutons. (**A**) Representative images showing dendritic spines in GCs *(left)* and mf boutons *(right),* and the associated line scan analysis of calcium transients (CaTs) elicited by uncaging of MNI-glutamate (see Methods), in control and *Grin1*-cKO animals. Blue dots indicate uncaging spots. Red Channel, Alexa 594; Green Channel, Fluo-5F. (**B**) Line scan analysis of CaTs measuring ΔG/R in dendritic spines when MNI-glutamate is uncaged in control or *Grin1*-cKO animals. Blue dots indicate location of two photon uncaging (2PU) pulses. (**C**) Summary plot demonstrating a significant reduction in dendritic spine CaTs in *Grin1*-cKO as compared to Control (control 0.053 ± 0.01 ΔG/R, n = 6 dendritic spines, 6 animals; *Grin1*-cKO 0.004 ± 0.003 ΔG/R, n = 6 spines, 6 animals; ΔG/R control vs. *Grin1*-cKO, p = 0.00088, unpaired *t*-test). (**D**) Line scan analysis of CaTs measuring ΔG/R in mf boutons when MNI-glutamate is uncaged in control or *Grin1*-cKO animals. (**E**) Summary plot demonstrating significant CaTs in boutons of control as compared to *Grin1*-cKO (control 0.014 ± 0.005, n = 6 boutons, 6 animals; *Grin1*-cKO -0.00012 ± -0.0006, n = 6 boutons, 6 animals; control vs. *Grin1*-cKO, p = 0.015, unpaired *t-*test). Data are presented as mean ± s.e.m. * p < 0.05; **** p < 0.001.

### PreNMDARs promote BDNF release from mossy fiber boutons

Previous work implicated preNMDARs in the release of BDNF at corticostriatal synapses following burst stimulation and presynaptic Ca^2+^ elevations (Park et al., 2014). Given the high expression levels of BDNF in mfs (Conner et al., 1997; Yan et al., 1997), we examined the potential role for preNMDARs in BDNF release from mf terminals. To this end, a Cre-dependent BDNF reporter (BDNF-pHluorin) was injected in *Grin1*-floxed and control animals. Littermate mice were injected with a mix of AAV5-CamKII-mCherry-Cre + AAV-DJ-DIO-BDNF-pHluorin viruses in the DG (**Figure 7A**). At least four weeks after surgery, acute slices were prepared for two-photon laser microscopy to image mf boutons. After acquiring a stable baseline of BDNF-pHluorin signals, mfs were repetitively activated (see Methods) (**Figure 7B**). BDNF-pHluorin signals were analyzed by measuring ΔF/F, where ΔF/F reductions indicate BDNF release (Park et al., 2014). We found that GluN1-deficient mf boutons showed a significant (∼50%) impairment in BDNF release as compared to control (**Figure 7C-D**). Furthermore, using a more physiological pattern of burst stimulation, GluN1-lacking mf boutons still displayed altered BDNF release as compared to control (**Figure 7-figure supplement 1**). Taken together, our results suggest preNMDARs contribute significantly to BDNF release during repetitive or burst stimulation of mf synapses.

**Figure 7.**
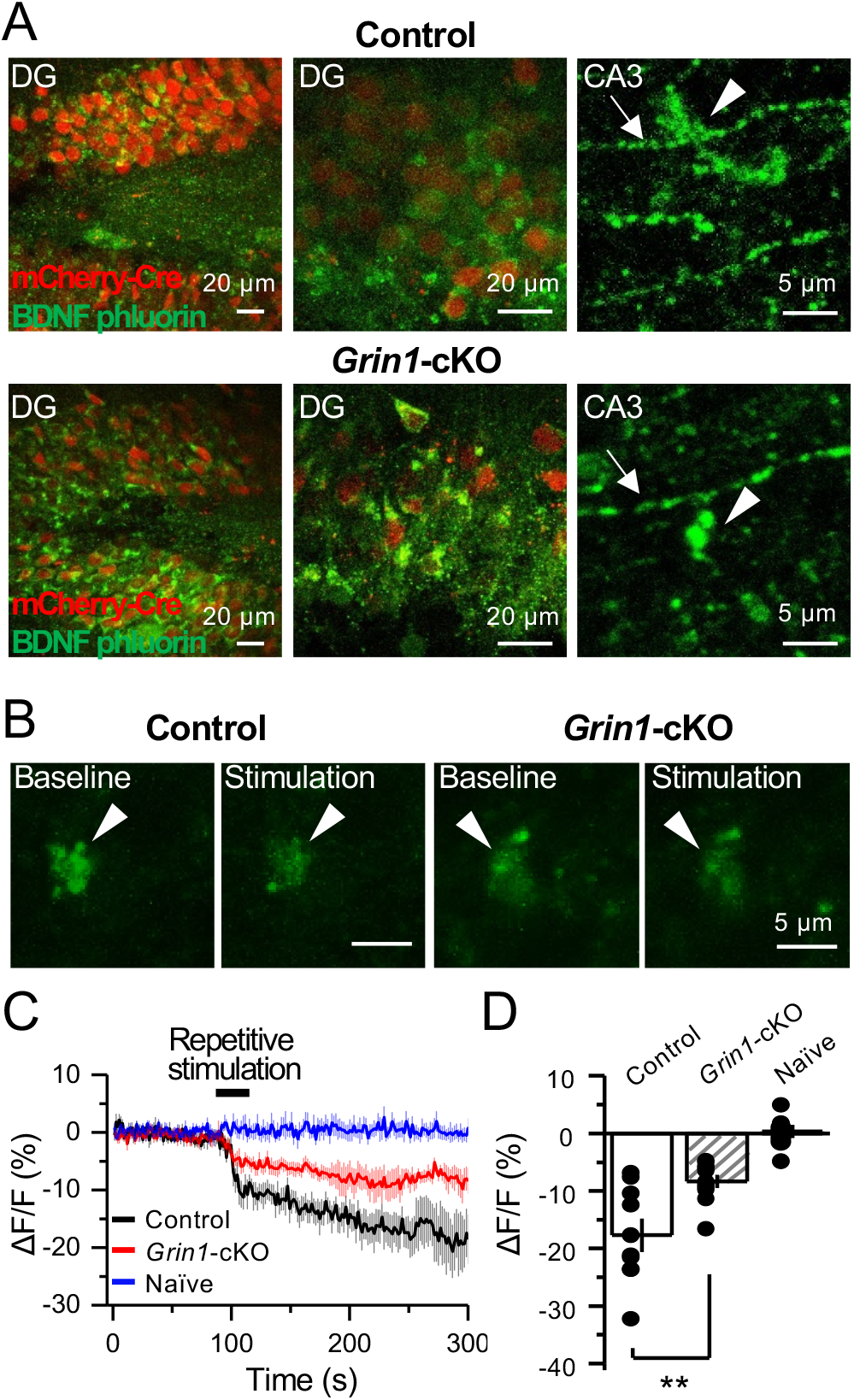
preNMDARs contribute significantly to BDNF release following repetitive activity. (**A**) Representative images showing expression of BDNF-pHluorin in the DG and CA3 area (arrows indicate mf axon, arrowheads indicate mf boutons). Control images *(top)*, *Grin1*- cKO images *(bottom)*. (**B**) Representative images of BDNF-pHluorin signal intensity at baseline and after repetitive stimulation of mfs (125 pulses, 25 Hz, x 2). Control images *(left)*, *Grin1*-cKO images *(right*), arrowhead indicates region of interest. (**C**) Time course of BDNF-pHluorin signal intensity measured as ΔF/F (%): control *(black)*, *Grin1*-cKO *(red),* Naïve *(blue)*. (**D**) Quantification of BDNF-pHluorin signal in (C) during the last 100 seconds reveals larger BDNF release in control animals as compared to *Grin1*-cKO (control -18% ± 3%, n = 9 slices, 5 animals; *Grin1*-cKO -8 ± 1%, n = 10 slices, 5 animals; *Grin1*-cKO vs. control, p = 0.00648, unpaired *t*-test). Data are presented as mean ± s.e.m. ** p < 0.01.

### PreNMDAR-mediated regulation of mossy fiber synapses is input-specific

In addition to providing a major excitatory input to the hippocampus proper, mf axons also synapse onto excitatory hilar mossy cells and inhibitory neurons in CA3 (Amaral et al., 2007; Henze et al., 2000; Lawrence & McBain, 2003). To test whether preNMDARs could also play a role at these synapses, we visually patched mossy cells and interneurons in acute rat hippocampal slices, loaded them with 35 µM Alexa 594 (**Figure 8A**) and 2 mM MK-801, and monitored AMPAR-EPSCs (V_h_ = -70 mV). Unlike mf-CA3 synapses, mf synapses onto CA3 interneurons in *stratum lucidum* do not express LFF, but can undergo burst-induced facilitation or depression (Toth et al., 2000). We found that MK-801 bath application had no effect on burst-induced facilitation or depression (**Figure 8B**), suggesting preNMDARs do not play a role at mf- Interneuron synapses in CA3. Mf inputs onto hilar mossy cells undergo robust activity-dependent facilitation (Lysetskiy et al., 2005). Similar to mf-CA3 synapses, we found that MK- 801 reduced LFF (**Figure 8C**). Stability experiments of mf transmission at CA3 interneurons or hilar mossy cells showed no significant differences (**Figure 8-figure supplement 1**). Taken together, our findings demonstrate that preNMDARs facilitate mf transmission onto excitatory neurons but not onto inhibitory interneurons.

**Figure 8.**
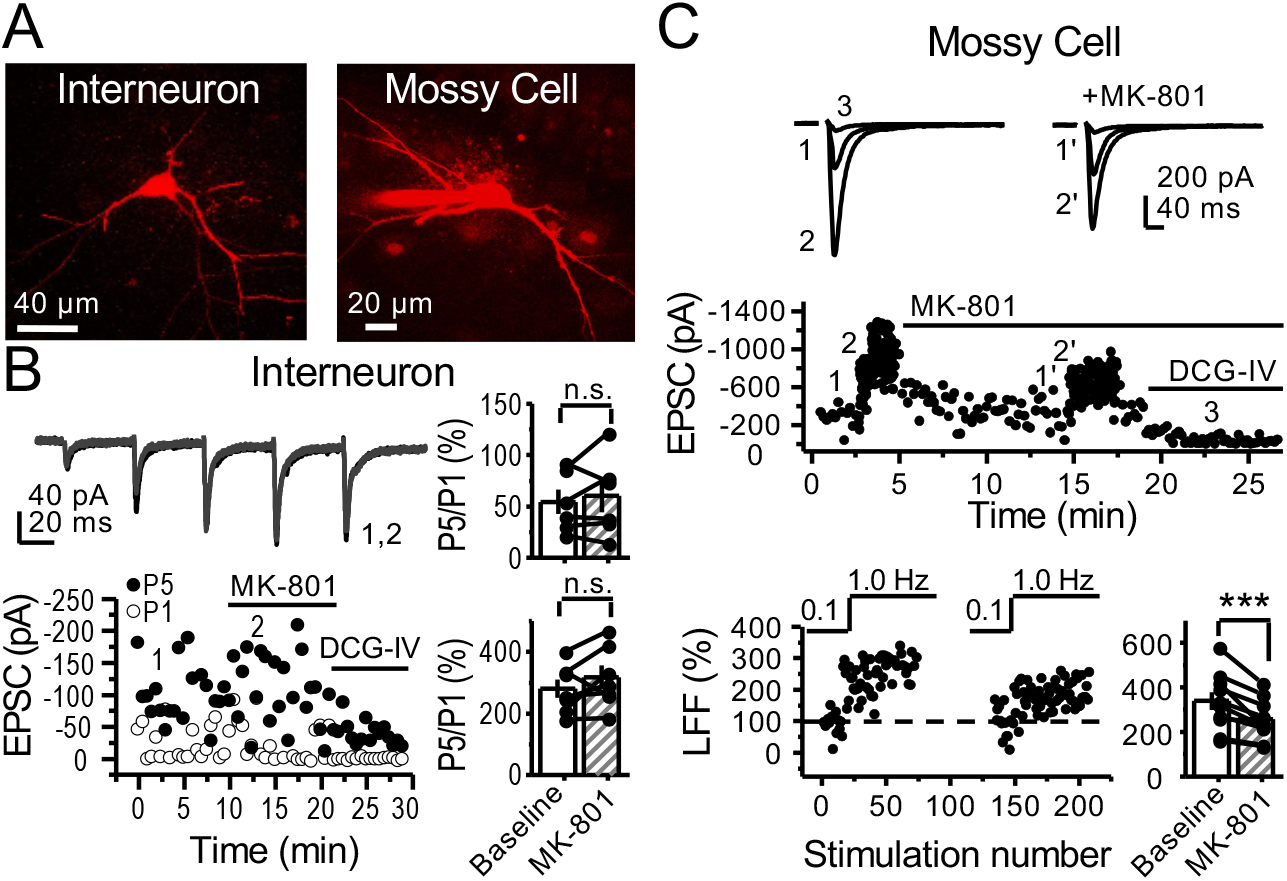
preNMDARs contribute to synaptic facilitation of mossy fiber inputs onto mossy cells but not onto CA3 inhibitory interneurons. (**A**) Representative images showing a CA3 interneuron and a hilar mossy cell patch-loaded with Alexa 594 (35 µM) for morphological identification in acute rat hippocampal slices. (**B**) AMPAR-EPSCs were recorded from CA3 interneurons at V_h_= -65 mV and burst-stimulation was elicited by 5 pulses at 25 Hz, see traces *(top)*. Representative experiment *(bottom, left)*, and summary plots *(right)* showing bath- application of MK-801 (50 µM) had no significant effect on depression *(top, right)* or facilitation *(bottom, right)* measured by P5/P1 ratio (baseline 54 ± 12%, MK-801 60 ± 16%, n = 6 cells; MK- 801 vs baseline, p = 0.675, Wilcoxon Signed Ranks test; baseline 281 ± 30%, MK-801 318 ± 37%, n = 7 cells; MK-801 vs baseline, p = 0.178, paired *t*-test, 5 animals in each data set). (**C**) AMPAR-ESPCs were recorded at V_h_ = -70 mV from hilar mossy cells, LFF was induced by stepping stimulation frequency from 0.1 to 1 Hz, see traces *(top)*. Representative experiment *(middle)*, normalized LFF and summary plot *(bottom)* indicating bath-application of MK-801 (50 µM) reduced facilitation (baseline 339 ± 41%, MK-801 258 ± 29%, n = 10 cells, 6 animals; baseline vs MK-801, p = 0.00152, paired *t*-test). DCG-IV (1 µM) was applied at the end of all experiments. Data are presented as mean ± s.e.m. *** p < 0.005.

## Discussion

In this study, we provide evidence that hippocampal mf boutons express preNMDARs whose activation fine-tunes mf synaptic function. Specifically, our results show that preNMDARs enhance mf short-term plasticity in a target cell-specific manner. By enhancing glutamate release onto excitatory but not inhibitory interneurons, preNMDARs increase GC-CA3 spike transfer. Moreover, using two-photon Ca^2+^ imaging, we demonstrate that preNMDARs contribute to presynaptic Ca^2+^ rise in mf boutons. Lastly, upon repetitive activity, preNMDARs promote BDNF release from mf boutons. Taken together, our findings indicate that preNMDARs act as autoreceptors to boost both glutamate and BDNF release at mf synapses. By regulating information flow in the DG-CA3 circuit, preNMDARs may play a significant role in learning and memory.

Early studies using immunoperoxidase electron microscopy revealed NMDARs in presynaptic compartments in multiple brain areas (for a review, see Corlew et al., 2008). Subsequent studies that used immunogold electron microscopy, a more precise localization method, identified NMDARs on the presynaptic membrane in a number of brain structures, including neocortex (Fujisawa & Aoki, 2003; Larsen et al., 2011), hippocampus (Berg et al., 2013; Jourdain et al., 2007; McGuinness et al., 2010), and amygdala (Pickel et al., 2006). In agreement with these studies, and using a previously validated antibody (Siegel et al., 1994), we identified prominent presynaptic labeling of the obligatory subunit GluN1 in mf boutons (Figure 1A-D). Moreover, we found that these receptors are close to the active zone and therefore well positioned to modulate neurotransmitter release.

Previous work in the cerebellum and neocortex suggested that somatodendritic potentials generated by NMDARs could signal to nerve terminals and lead to presynaptic Ca^2+^ elevations (Christie & Jahr, 2008, 2009). Thus, changes in neurotransmitter release resulting from NMDAR antagonism could be due to somatodendritic NMDARs but not necessarily preNMDARs residing on nerve terminals (Duguid, 2013). However, we showed that focal NMDAR antagonism far from the somatodendritic compartment and in transected axons still reduced short-term plasticity at mf synapses (Figure 3), making it extremely unlikely that somatodendritic NMDARs could explain our results. In further support of functional preNMDARs at mf boutons, we found that 2PU of glutamate induced Ca^2+^ rise in control but not in GluN1-deficient boutons. Together, our findings strongly support the presence of functional preNMDARs facilitating neurotransmission at mf-CA3 synapses.

There is evidence that preNMDARs can operate as coincidence detectors at some synapses (Duguid, 2013; Wong et al., 2020). At the mf-CA3 synapse, we found that preNMDARs contribute to LFF (i.e. 1s inter-stimulus interval). This observation is intriguing given that the presynaptic AP-mediated depolarization is likely absent by the time glutamate binds to preNMDARs. However, coincidence detection may not be an essential requirement for mf preNMDARs to modulate glutamate release. Of note, at resting membrane potential the NMDAR conductance is not zero and the driving force for Ca^2+^ influx is high (Paoletti et al., 2013; Traynelis et al., 2010). It is also conceivable that mf preNMDARs exhibit low-voltage dependence, as it has been reported at other synapses (Wong et al., 2020). Remarkably, the somatodendritic compartment of GCs can generate sub-threshold depolarizations at mf terminals (a.k.a. excitatory presynaptic potentials) (Alle & Geiger, 2006). By alleviating the magnesium blockade, these potentials might reduce the need for coincidence detection and transiently boost the functional impact of mf preNMDARs.

While the presence of preNMDARs is downregulated during development both in neocortex (Corlew et al., 2007; Larsen et al., 2011) and hippocampus (Mameli et al., 2005), we were able to detect functional preNMDARs in young adult rats (P17-P28) and mice (P30-P44), once mf connections are fully developed (Amaral & Dent, 1981). Functional preNMDARs have been identified in axonal growth cones of hippocampal and neocortical neurons, suggesting these receptors are important for regulating early synapse formation (Gill et al., 2015; Wang et al., 2011). Because GCs undergo adult neurogenesis, and adult born GCs establish new connections in the mature brain, preNMDARs could also play an important role at immature mf synapses and functional integration of new born GCs into the mature hippocampus (Toni & Schinder, 2015). Moreover, experience can modulate the expression and composition of preNMDARs in neocortex (Larsen et al., 2014), a possibility not investigated in our study.

The glutamate that activates preNMDARs may originate from the presynaptic terminal, the postsynaptic cell, nearby synapses or neighboring glial cells. Our results indicate that activation of preNMDARs at mf synapses requires activity-dependent release of glutamate that likely arises from mf boutons, although other sources cannot be discarded, including astrocytes. For instance, at medial entorhinal inputs to GCs, preNMDARs appear to be localized away from the presynaptic release sites and facing astrocytes, consistent with preNMDAR activation by gliotransmitters (Jourdain et al., 2007; Savtchouk et al., 2019). In contrast, at mf-CA3 synapses we found that preNMDARs are adjacent to the release sites suggesting a direct control on glutamate release from mf boutons.

The precise mechanism by which preNMDARs facilitate neurotransmitter release is poorly understood but it may include Ca^2+^ influx through the receptor and depolarization of the presynaptic terminal with subsequent activation of voltage-gated Ca^2+^ channels (Banerjee et al., 2016; Corlew et al., 2008). In support of this mechanism is the high Ca^2+^ permeability of NMDARs (Paoletti et al., 2013; Rogers & Dani, 1995). Besides, presynaptic subthreshold depolarization and subsequent activation of presynaptic voltage-gated Ca^2+^ channels is a common mechanism by which presynaptic ionotropic receptors facilitate neurotransmitter release (Engelman & MacDermott, 2004; Pinheiro & Mulle, 2008). PreNMDARs may also act in a metabotropic manner (Dore et al., 2016) and facilitate spontaneous transmitter release independent of Ca^2+^-influx (Abrahamsson et al., 2017). Our findings demonstrating that the open channel blocker MK-801 robustly reduced short-term plasticity at mf synapses support an ionotropic mechanism that involves Ca^2+^ influx through preNMDARs. A previous study failed to observe Ca^+2^ reductions in mf boutons by DL-APV (Liang et al., 2002). A combination of factors could account for this discrepancy with our study, including a stronger mf repetitive stimulation (20 pulses, 100 Hz) which may overcome a less potent NMDAR antagonism and/or the need for preNMDAR activity, as well as the use of a higher affinity Ca^2+^ indicator (Fura-2 AM) and a lower spatiotemporal resolution imaging approach. Nevertheless, in line with previous studies that detected presynaptic Ca^2+^ rises following local activation of NMDARs (e.g. NMDA or glutamate uncaging) in visual cortex (Buchanan et al., 2012) and cerebellum (Rossi et al., 2012), we provide direct evidence that preNMDAR activation either by repetitive activation of mfs or 2PU of glutamate increases presynaptic Ca^2+^ (Figures 5 and 6). Although the Ca^2+^ targets remain unidentified, these may include proteins of the release machinery, calcium-dependent protein kinases and phosphatases, and Ca^2+^ release from internal stores (Banerjee et al., 2016). In addition to facilitating evoked neurotransmitter release, preNMDARs can promote spontaneous neurotransmitter release as indicated by changes in miniature, action potential-independent activity (e.g. mEPSCs) (for recent reviews, see Banerjee et al., 2016; Kunz et al., 2013; Wong et al., 2020). A potential role for preNMDARs in spontaneous, action potential-independent release at mf synapses cannot be discarded.

Our results show that activation of preNMDARs by physiologically relevant patterns of presynaptic activity enhanced mf transmission and DG-CA3 information transfer (Figure 4). A previous study reported NMDAR genetic deletion in GCs resulted in memory deficits (e.g. pattern separation) (McHugh et al., 2007). Although the mechanism is unclear, it could involve activity-dependent preNMDAR regulation of mf excitatory connections. We also found that preNMDARs facilitate neurotransmitter release in a target cell-specific manner. Like in neocortex (Larsen & Sjostrom, 2015), such specificity strongly suggests that preNMDARs have distinct roles in controlling information flow in cortical microcircuits. Thus, preNMDAR facilitation of mf synapses onto glutamatergic neurons but not GABAergic interneurons (Figure 8) may fine-tune the CA3 circuit by increasing the excitatory/inhibitory balance.

Given the multiple signaling cascades known to regulate NMDARs (Lau & Zukin, 2007; Sanz-Clemente et al., 2013), preNMDARs at mf synapses may provide an important site of neuromodulatory control. PreNMDARs have been implicated in the induction of LTP and LTD at excitatory or inhibitory synapses in several brain areas (Banerjee et al., 2016; Wong et al., 2020). While most evidence, at least using robust induction protocols *in vitro,* indicates that long-term forms of presynaptic plasticity at mf synapses can occur in the absence of NMDAR activation (Castillo, 2012; Nicoll & Schmitz, 2005), our findings do not discard the possibility that preNMDARs could play a role *in vivo* during subtle presynaptic activities. As previously reported for corticostriatal LTP (Park et al., 2014), preNMDARs could regulate long-term synaptic plasticity by controlling BDNF release, which is consistent with BDNF-TrkB signaling being implicated in mf-CA3 LTP (Schildt et al., 2013). In addition, BDNF could facilitate glutamate release by enhancing NMDAR function at the presynapse, as previously suggested (W. Chen et al., 2014; Madara & Levine, 2008), although the precise mechanism(s) remain unclear. By potentiating mf-CA3 transmission, BDNF could also promote epileptic activity (McNamara & Scharfman, 2012). Lastly, dysregulation of NMDARs is commonly implicated in the pathophysiology of brain disorders such as schizophrenia, autism, and epilepsy (Lau & Zukin, 2007; Paoletti et al., 2013). PreNMDAR expression and function have been suggested to be altered in experimental models of disease, including neuropathic pain (Y. Chen et al., 2019; Zeng et al., 2006), and epilepsy (Yang et al., 2006). At present, however, *in vivo* evidence for the involvement of preNMDARs in brain function and disease is rather indirect (Bouvier et al., 2015; Wong et al., 2020). The development of specific preNMDAR tools is required to determine the functional impact of these receptors *in vivo*.

## Methods

### Antibodies

A monoclonal antibody against GluN1 (clone 54.1 MAB363) was obtained from Millipore (Germany) and its specificity was characterized previously (Siegel et al., 1994). An affinity-purified polyclonal rabbit anti-GluA1-4 (pan-AMPA), corresponding to aa 724–781 of rat, was used and characterised previously (Nusser et al., 1998).

### Immunohistochemistry for electron microscopy

Immunohistochemical reactions at the electron microscopic level were carried out using the post-embedding immunogold method as described earlier (Lujan et al., 1996). Briefly, animals (n = 3 rats were anesthetized by intraperitoneal injection of ketamine-xylazine 1 : 1 (0.1 mL/kg b.w.) and transcardially perfused with ice-cold fixative containing 4% paraformaldehyde, 0.1% glutaraldehyde and 15% saturated picric acid solution in 0.1 M phosphate buffer (PB) for 15 min. Vibratome sections 500 μm thick were placed into 1 M sucrose solution in 0.1 M PB for 2 h before they were slammed on a Leica EM CPC apparatus. Samples were dehydrated in methanol at -80°C and embedded by freeze-substitution (Leica EM AFS2) in Lowicryl HM 20 (Electron Microscopy Science, Hatfield, USA), followed by polymerization with UV light. Then, ultrathin 80-nm-thick sections from Lowicryl-embedded blocks of the hippocampus were picked up on coated nickel grids and incubated on drops of a blocking solution consisting of 2% human serum albumin in 0.05 M TBS and 0.03% Triton X-100. The grids were incubated with GluN1 or pan-AMPA antibodies (10 μg/mL in 0.05 M TBS and 0.03% Triton X-100 with 2% human serum albumin) at 28 °C overnight. The grids were incubated on drops of goat anti-rabbit IgG conjugated to 10 nm colloidal gold particles (Nanoprobes Inc.) in 2% human serum albumin and 0.5% polyethylene glycol in 0.05 M TBS and 0.03% Triton X-100. The grids were then washed in TBS and counterstained for electron microscopy with 1% aqueous uranyl acetate followed by Reynolds’s lead citrate. Ultrastructural analyses were performed in a JEOL-1010 electron microscope.

### Hippocampal slice preparation

Animal handling followed an approved protocol by the Albert Einstein College of Medicine Institutional Animal Care and Use Committee in accordance with the National Institute of Health guidelines. Acute rat hippocampal slices (400 µm thick) were obtained from Sprague-Dawley rats, from postnatal day 17 (P17) to P28 of either sex. For procedures regarding transgenic mouse slice preparation, see below. The hippocampi were isolated and cut using a VT1200s microslicer (Leica Microsystems Co.) in a solution containing (in mM): 215 sucrose, 2.5 KCl, 26 NaHCO_3_, 1.6 NaH_2_PO_4_, 1 CaCl_2_, 4 MgCl_2_, 4 MgSO_4_ and 20 glucose. Acute slices were placed in a chamber containing a 1:1 mix of sucrose cutting solution and normal extracellular artificial cerebrospinal fluid (ACSF) recording solution containing (in mM): 124 NaCl, 2.5 KCl, 26 NaHCO_3_, 1 NaH_2_PO_4_, 2.5 CaCl_2_, 1.3 MgSO_4_ and 10 glucose incubated in a warm-water bath at 33-34 °C. The chamber was brought to room temperature for at least 15 min post-sectioning and the 1:1 sucrose-ACSF solution was replaced by ACSF. All solutions were equilibrated with 95% O_2_ and 5% CO_2_ (pH 7.4). Slices were allowed to recover for at least 45 min in the ACSF solution before recording. For physiological Mg^+2^ and Ca^+2^ experiments, ACSF solutions were adjusted to (in mM): 1.2 MgSO_4_, and 1.2 CaCl_2_ and temperature was maintained at 35 ± 0.1 °C in the submersion-type recording chamber heated by a temperature controller (TC-344B Dual Automatic Temperature Controller, Warner Instruments).

### Electrophysiology

Electrophysiological recordings were performed at 26.0 ± 0.1 °C (unless otherwise stated) in a submersion-type recording chamber perfused at 2 mL/min with normal ACSF supplemented with the GABA_A_ receptor antagonist picrotoxin (100 µM) and the selective AMPA receptor (AMPAR) antagonist LY303070 at a low concentration (0.5 µM) to minimize CA3-CA3 recurrent activity, or at a high concentration (15 µM) to isolate KAR-EPSCs and KAR-EPSPs to assess monosynaptic mf transmission. Whole-cell recordings were made from CA3 pyramidal cells voltage clamped at -70 mV using patch-type pipette electrodes (3-4 mΩ) containing (in mM): 131 cesium gluconate, 8 NaCl, 1 CaCl_2_, 10 EGTA, 10 glucose, 10 HEPES, and 2 MK-801 pH 7.25 (280-285 mOsm) unless specified otherwise. KOH was used to adjust pH. Series resistance (8-15 MΩ) was monitored throughout all experiments with a -5 mV, 80 ms voltage step, and cells that exhibited a series resistance change (>20%) were excluded from analysis. A stimulating bipolar electrode (theta glass, Warner Instruments) was filled with ACSF and placed in *stratum lucidum* to selectively activate mfs using a DS2A Isolated Voltage Stimulator (Digitimer Ltd.) with a 100 µs pulse width duration. AMPAR-EPSCs were recorded for a baseline period of two minutes and low-frequency facilitation (LFF) was induced by stepping the stimulation frequency from 0.1 to 1 Hz for two minutes. Facilitation was measured by taking a ratio of the mean EPSC during the steady-state, LFF period of activity and the two-minute baseline (EPSC_1Hz_/EPSC_0.1Hz_) before and after bath-application of NMDAR antagonists.

To qualify for analysis, mf responses met three criteria: 1) The 20-80% rise time of the AMPAR-EPSC was less than 1 ms, 2) LFF was greater than 150%, 3) The AMPAR-EPSC displayed at least 70% sensitivity to the group 2/3 mGluR agonist, DCG-IV (1 µM). Isolated KAR-EPSCs were elicited by 2 pulses with a 5 ms inter-stimulus-interval for LFF experiments. Baseline measurements were acquired at least 10 min after “break-in” to achieve optimal intracellular blockade of postsynaptic NMDARs by MK-801 (2 mM) in the patch-pipette. To transect mf axons in acute slices, a 45° ophthalmic knife (Alcon Surgical) was used to make a diagonal cut across the hilus from the dorsal to ventral blades of the DG, and the subregion CA3b was targeted for patch-clamp recordings. For D-APV (2 mM) puff experiments, a puffer device (Toohey Company) was set to deliver 2-3 puffs of 100 ms duration at 3-4 psi during the two minutes of LFF activity. The puffer pipette was placed at least 200 µm away from the recording site and both the puff pipette and hippocampal slice were positioned to follow the direction of the laminar perfusion flow in a low profile, submersion-type chamber (RC-26GLP, Warner Instruments). Burst-induced facilitation was elicited by 5 pulses at 25 Hz with a 0.03 Hz inter-trial-interval for a baseline period of 10 min. Facilitation was measured by calculating the ratio of the mean KAR-EPSC peak of the 5^th^ pulse to the 1^st^ pulse (P_5_/P_1_) before and after bath-application of MK-801 (50 µM). To study KAR induced action potentials, CA3 pyramidal cells were whole-cell patch-clamped with internal solution containing in (mM): 112 potassium gluconate, 17 KCl, 0.04 CaCl_2_, 0.1 EGTA, 10 HEPES, 10 NaCl, 2 MgATP, 0.2 Na_3_GTP and 2 MK-801, pH 7.2 (280-285 mOsm). Current-clamped CA3 cells were held at -70 mV during burst stimulation of mfs (5 pulses at 25 Hz) to monitor evoked action potentials. Spike-transfer was measured by quantifying mean number of spikes/burst for a 10 min period before and after bath application of MK-801 (50 µM). Robust sensitivity to the AMPAR/KAR selective antagonist NBQX (10 µM) confirmed KAR-EPSC responses. Similarly, CA3 pyramidal cells were kept in current-clamp mode for AMPAR-mediated action potential monitoring in the presence of LY303070 (0.5 µM) and picrotoxin (100 µM). AMPAR-mediated mf action potentials were confirmed by blockade of responses following application of DCG-IV (1 µM). Both hilar mossy cells and CA3 interneurons were visually patched-loaded with Alexa 594 (35 µM) and morphological identity was confirmed by two-photon laser scanning imaging at the end of experiments. Hilar mossy cells were voltage clamped at -70 mV and a bipolar electrode was placed in the DG to activate mf inputs. The data analysis and inclusion criteria used for mf experiments (described above) was also implemented for hilar mossy cell recordings. CA3 interneurons were voltage clamped at -70 mV and burst-stimulated, facilitation was assessed as previously mentioned. Both facilitating and depressing mf responses were included for analysis given the diversity of mf-CA3 interneuron transmission (Toth et al., 2000). Whole-cell voltage and current clamp recordings were performed using an Axon MultiClamp 700B amplifier (Molecular Devices). Signals were filtered at 2 kHz and digitized at 5 kHz. Stimulation and acquisition were controlled with custom software (Igor Pro 7).

### Transgenic animals

*Grin1*-floxed littermate mice of either sex (P16-20) were injected with 1 μL of AAV5-CamKII-eGFP, AAV5-CamKII-CreGFP, AAV5-CamKII-mCherry, or AAV5-CamKII-mCherry-Cre viruses at a rate of 0.12 μL/min at coordinates (-1.9 mm A/P, 1.1 mm M/L, 2.4 mm D/V) targeting the DG using a stereotaxic apparatus (Kopf Instruments). Two weeks post-surgery mice were sacrificed for electrophysiology or Ca^2+^ imaging experiments. Mice were transcardially perfused with 20 mL of cold NMDG solution containing in (mM): 93 NMDG, 2.5 KCl, 1.25 NaH_2_PO_4_, 30 NaHCO_3_, 20 HEPES, 25 glucose, 5 sodium ascorbate, 2 Thiourea, 3 sodium pyruvate, 10 MgCl_2_, 0.5 CaCl_2_, brought to pH 7.35 with HCl. The hippocampi were isolated and cut using a VT1200s microslicer in cold NMDG solution. Acute mouse slices were placed in an incubation chamber containing normal ACSF solution that was kept in a warm water bath at 33-34 °C. All solutions were equilibrated with 95% O_2_ and 5% CO_2_ (pH 7.4). Post-sectioning, slices recovered at room temperature for at least 45 min prior to experiments. For NMDAR/AMPAR ratios, GCs were patch-clamped with the cesium internal solution previously mentioned containing either Alexa 594 (35 µM) for GFP^+^ cells (laser tuned to 830 nm/910 nm, respectively) or Alexa 488 (35 µM) for mCherry^+^ cells (laser tuned to 910 nm/780 nm, respectively). AMPAR-EPSCs were recorded at -65 mV in the presence of picrotoxin (100 µM) by placing a bipolar electrode near the medial perforant path and delivering a 100 μs pulse width duration using an Isoflex stimulating unit. AMPAR-EPSCs were acquired for at least 5 min followed by bath-application of NBQX (10 µM) to isolate NMDAR-EPSCs. GCs were brought to +40 mV to alleviate magnesium block and record optimal NMDAR-EPSCs. NMDAR/AMPAR ratios were measured by taking the mean NMDAR-EPSC/AMPAR-EPSC for a 5 min period of each component. Only acute mouse slices with optimal GFP and mCherry reporter fluorescence (i.e. robust expression, ≥ 75% of dentate gyrus fluorescence) were used for electrophysiology, and Ca^2+^ and BDNF imaging experiments. *Grin1*-floxed animals (The Jackson Laboratory) were kindly provided by Dr. Michael Higley (Yale University).

### Optogenetics

*Grin1* floxed and control mice of either sexes (P17-20) were injected with a 1:2 mix of AAV5- CamKII-CreGFP/AAV-DJ-FLEX-ChIEF-tdTomato viruses targeting the DG, using the same coordinates described above. At least four weeks post-surgery acute hippocampal slices were prepared as previously described and slices showing optimal GFP and tdTomato expression were used for electrophysiology experiments. Mf optical burst-stimulation was elicited by using a Coherent 473 nm laser (4-8 mW) delivering 5 pulses at 25 Hz with a 1-2 ms pulse width duration. Facilitation was measured by taking a ratio of the mean AMPAR-EPSC peak of the 5^th^ pulse to the 1^st^ pulse (P_5_/P_1_) in control and *Grin1*-cKO animals.

### Two-photon calcium imaging and MNI-glutamate uncaging

mCherry+ GCs were patch-loaded with an internal solution containing in (mM): 130 KMeSO_4_, 10 HEPES, 4 MgCl_2_, 4 Na_2_ATP, 0.4 NaGTP, 10 sodium phosphocreatine, 0.035 Alexa 594 (red morphological dye), and 0.2 Fluo-5F (green calcium indicator), 280-285 mOsm. KOH was used to adjust pH. GCs near the hilar border were avoided and GCs that exhibited adult-born GCs electrophysiological properties were excluded from analysis. GCs were kept in voltage clamp configuration at -50 mV for at least 1 hr to allow the diffusion of dyes to mf boutons. Recordings were obtained in ACSF solution containing in (mM): 124 NaCl, 2.5 KCl, 26 NaHCO_3_, 1 NaH_2_PO_4_, 4 CaCl_2_, 0 MgSO_4_, 10 glucose, 0.01 NBQX, 0.1 picrotoxin, and 0.01 D-serine. Using an Ultima 2P laser scanning microscope (Bruker Corp) equipped with an Insight Deep See laser (Spectra Physics) tuned to 830 nm, the "red" photomultiplier tube (PMT) was turned on and with minimal pockel power the red signal was used to identify the mf axon. With 512 x 512 pixel resolution mf axons were followed for at least 200 µm from the DG towards CA3, until bouton structures were morphologically identified and measured (>3 μm in diameter). GCs were switched to current clamp mode held at -70 mV and 1 ms current injections were used to elicit a burst of 5 action potentials at 25 Hz. Using line scan analysis software (PrairieView 5.4, Bruker Corp.), a line was drawn across the diameter of the bouton at a magnification of at least 16X. The “green” PMT channel was turned on and 1,000 line scans were acquired in a 2 s period. Action potential induction was delayed for 400 ms to collect a baseline fluorescence time period. Calcium transients (CaTs) were acquired with a 1 min inter-trial-interval and analyzed using the ΔG/R calculation: (G - G_0_)/R. CaTs from control animals were compared to *Grin1-*cKO by taking the mean peak ΔG/R value for a 30 ms period of the 5^th^ action potential. In similar fashion, CaT signals in acute rat hippocampal slices were acquired and tested for sensitivity to D-APV (100 µM) while adjusting ACSF MgS0_4_ concentration to 1.3 mM and CaCl_2_ to 2.5 mM in the absence of NBQX.

For glutamate uncaging experiments, GCs that were mCherry+ were patch-loaded using the internal solution previously described, and a small volume (12 mL) of recirculated ACSF solution containing in (mM): 124 NaCl, 2.5 KCl, 26 NaHCO_3_, 1 NaH_2_PO_4_, 4 CaCl_2_, 0 MgSO_4_, 10 glucose, 2.5 MNI-glutamate, 0.01 NBQX, 0.1 picrotoxin, and 0.01 D-serine. A MaiTai HP laser (Spectra Physics) was tuned to 720 nm to optimally uncage glutamate and elicit CaTs in GC dendritic spines. Following the measurement of CaTs in GC spines, mf boutons were identified and to mimic bursting activity, 5 uncaging pulses (1 ms duration) were delivered at 25 Hz. The acquired CaTs in spines and boutons were analyzed using the ΔG/R calculation in control and *Grin1*-cKO animals. In a subset of control boutons, D-APV (100 µM) was applied to detect CaT sensitivity to NMDAR antagonism.

### Two-photon BDNF-phluorin imaging

*Grin1* floxed and control mice of both sexes (P16-20) were injected with a 1:2 mix of AAV5- CamKII-mCherryCre/AAV-DJ-DIO-BDNF-phluorin viruses targeting the DG using the same coordinates as above. At least four weeks post-surgery acute hippocampal slices were prepared as previously described and slices showing optimal GFP and mCherry expression were taken for imaging sessions. For stimulation, a monopolar micropipette electrode was placed in the *stratum lucidum* at least 250 µm away from the imaging site. The Insight Deep See laser (Spectra Physics) was tuned to 880 nm and the imaging site was selected by the appearance of fibers and bouton structures in the *stratum lucidum*. Using 512 X 512 pixel resolution identified boutons measuring at least 3 μm in diameter were selected as a region of interest (ROI) magnified to 4-6X and a baseline acquisition of 100 consecutive images at 1 Hz using T-series software (PrairieView 5.4, Bruker Corp.) was acquired (Park et al., 2014). Following the baseline acquisition a repetitive stimulation consisting of 125 pulses at 25 Hz was delivered 2x, triggering an acquisition of 200 consecutive images at 1 Hz. The fluorescence intensity of the bouton ROI was measured using ImageJ software to calculate ΔF/F of the BDNF-pHluorin signal. To verify reactivity of the ROI an isosmotic solution of NH_4_Cl (50 mM) was added at the end of the imaging session as previously reported (Park et al., 2014). The same experimental and analysis procedure was implemented to measure BDNF release triggered by mf burst stimulation consisting of 5 pulses at 100 Hz, 50x, every 0.5 s.

### Viruses

AAV5-CamKII-eGFP and AAV5-CamKII-CreGFP viruses were acquired from UPenn Vector Core. AAV5-CamKII-mCherry and AAV5-CamKII-mCherry-Cre were obtained from UNC Chapel Hill Vector Core. The AAV-DJ-FLEX-ChIEF-tdTomato and AAV-DJ-DIO-BDNF-phluorin viruses were custom ordered and obtained from UNC Chapel Hill Vector Core. The DNA of the ChIEF virus was a generous gift from Dr. Pascal Kaeser (Harvard University), and the DNA of the BDNF-pHluorin was kindly provided by Dr. Hyungju Park (Korea Brain Research Institute).

### Chemicals & Drugs

Picrotoxin and all chemicals used to prepare cutting, recording, and internal solutions were acquired from MilliporeSigma. All NMDAR antagonists (D-APV, MK-801, R-CPP), NMDAR agonist (D-serine), the group 2/3 mGluR agonist (DCG-IV) and MNI-glutamate for uncaging experiments were purchased from Tocris Bioscience. D-APV was also acquired from the NIMH Chemical Synthesis Drug Program. NBQX was purchased from Cayman Chemical Company. The noncompetitive AMPAR selective antagonist LY303070 was custom ordered from ABX Chemical Company. Alexa 594 morphological dye, Alexa 488, and the Ca^2+^ indicator Fluo-5F (Invitrogen) were purchased from ThermoFisher Scientific.

### Statistical analysis and Data Acquisition

All data points from experiments were tested for normality using a Shapiro-Wilk test (p value < 5% for a normal distribution). Statistical significance was determined if p value < 0.05. Experiments with a normal distribution and an N > to 7 cells were tested for statistical significance with a paired Student *t*-test. Experiments with N < 7 cells or skewed distributions were tested for statistical significance using a paired Wilcoxon signed rank sum test. For experiments comparing control and *Grin1*-cKO animals, statistical significance was determined using Unpaired *t*-test and Mann-Whitney test (U < 0.05). All statistical tests were performed using Origin Pro 9 (Origin Lab). Experimenters were blind to the identity of the virus injected in transgenic *Grin1* floxed mice during the acquisition of data in CA3 electrophysiology and two-photon imaging. However, data analysis could not be performed blind in those experiments in which NMDAR/AMPAR ratios in GCs were examined in order to assess the efficiency of the cKO.

---

**Figure 1-figure supplement 1.**
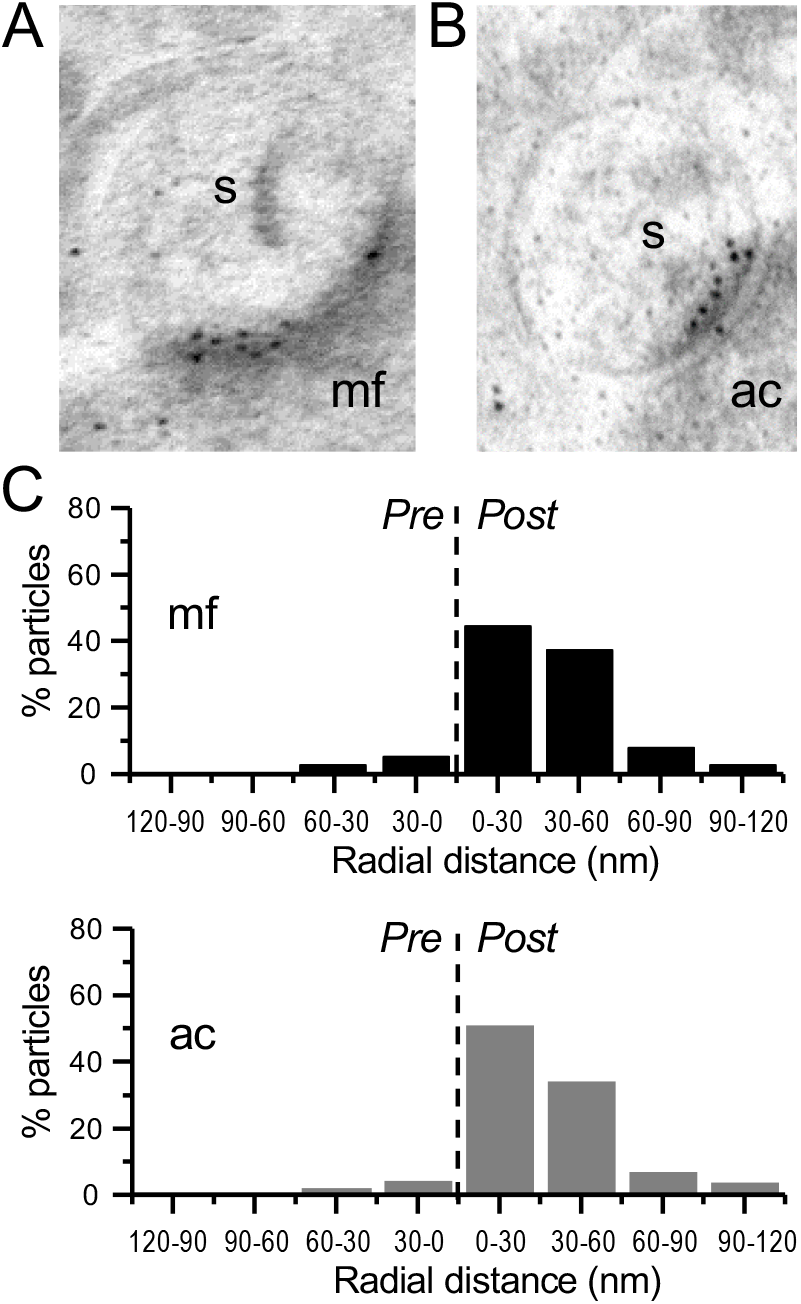
Immunogold-EM reveals negligible presynaptic AMPAR particle distribution. (**A,B**) Images of mossy fiber (mf) and associational commissural (ac) synapses, postsynaptic spines (s). (**C**) AMPAR immuno-particle distribution (30 nm bins), mf: 102 synapses, 8 presynaptic particles; ac: 75 synapses, 6 presynaptic particles; 3 animals. Dashed line represents synaptic cleft.

**Figure 1-figure supplement 2.**
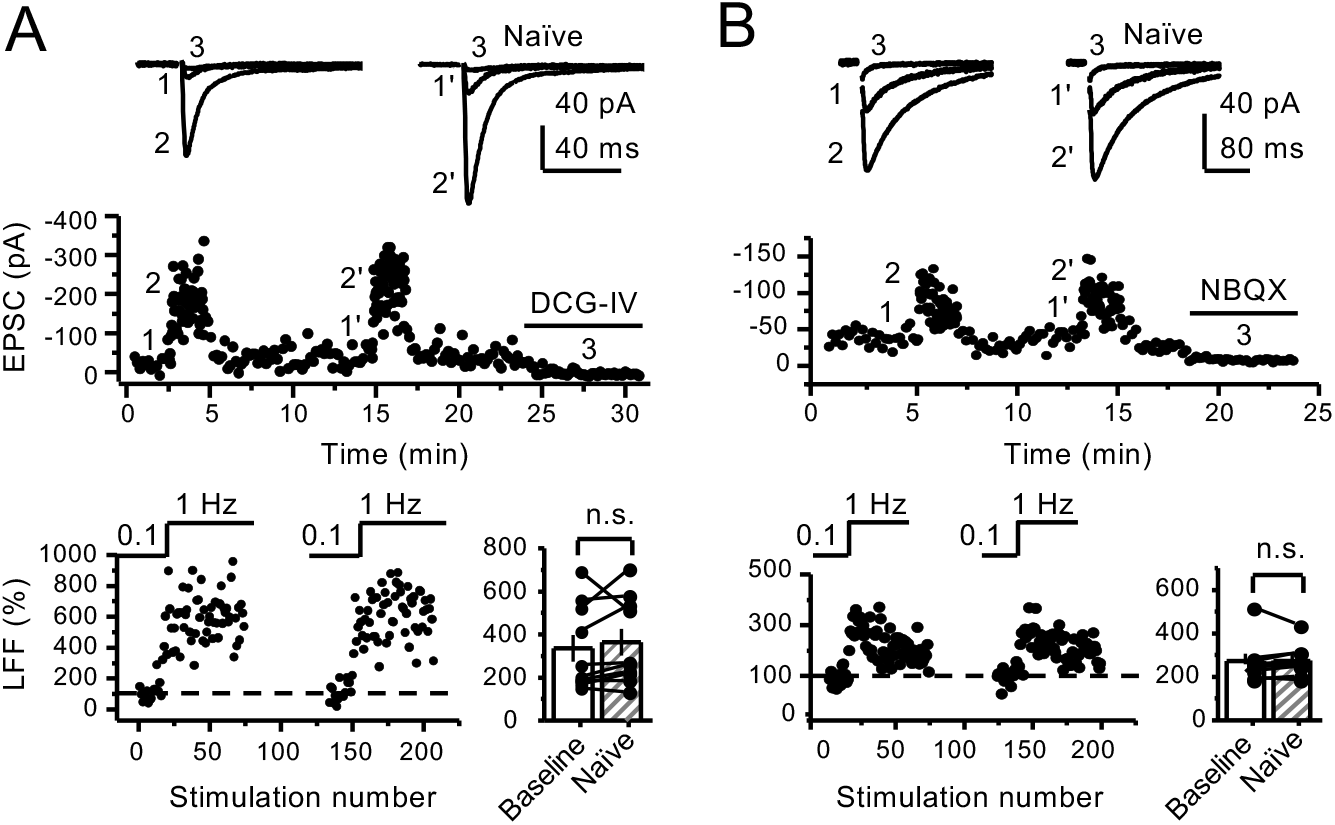
(**A**) Stable low-frequency facilitation (LFF) of AMPAR-EPSCs. In naïve slices (interleaved experiments), LFF remained unchanged throughout the recording session (baseline 335 ± 62%, naïve 363 ± 63%, n = 10 cells, 9 animals; p = 0.185, Wilcoxon-Signed Ranks test, baseline vs naÏve). DCG-IV (1 µM) was applied at the end of all recordings to confirm mf-CA3 transmission. (**B**) LFF of KAR-EPSCs was also stable in interleaved, naïve slices (baseline 274 ± 33%, naïve 278 ± 25%, n = 9 cells, 6 animals; p = 0.236, Wilcoxon Signed Ranks test, baseline vs naïve). NBQX (10 µM) was applied at the end of all recordings to confirm mf KAR transmission. Data are presented as mean ± s.e.m.

**Figure 1-figure supplement 3.**
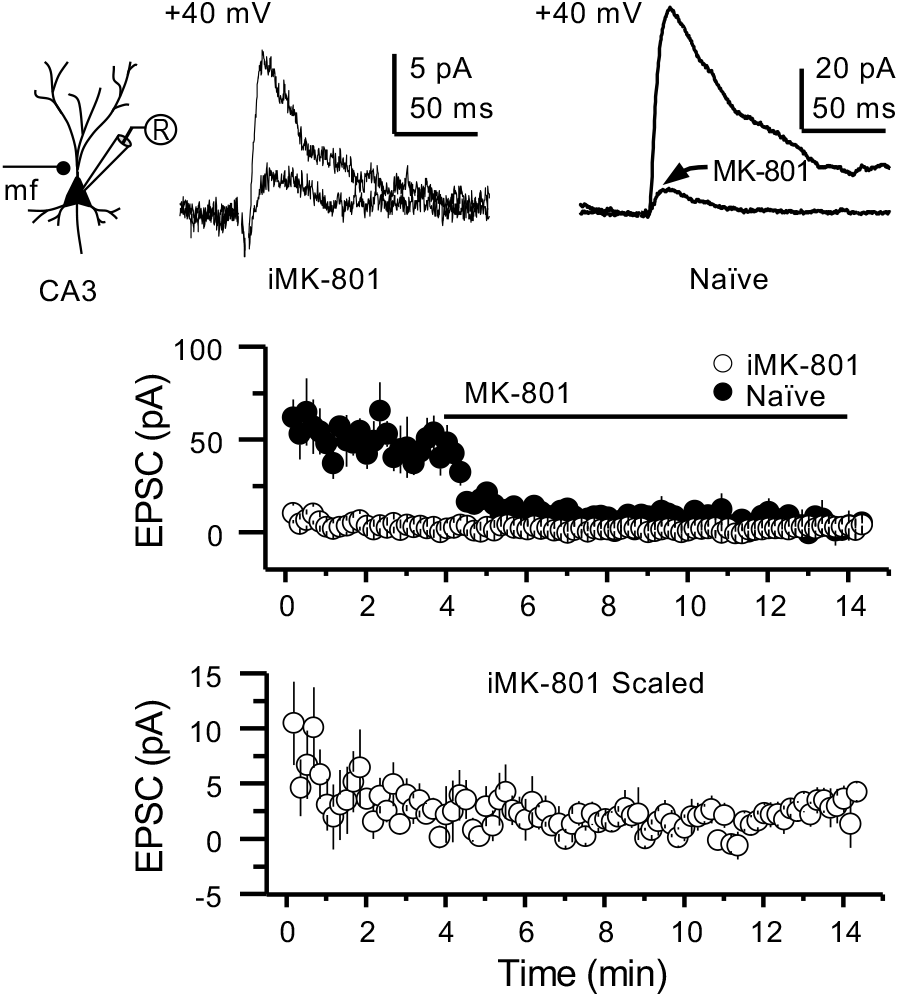
Intracellular MK-801 effectively blocked postsynaptic NMDARs. Representative NMDAR-EPSCs (V_h_ = + 40 mV) from CA3 pyramidal neurons patch-loaded with 2 mM MK-801 *(left)* or naïve internal solution *(right)*. Mf inputs were stimulated with a bipolar electrode (theta-glass pipette) in *stratum lucidum* delivering in the presence of picrotoxin (100 µM) and NBQX (10 µM). Bath-application of MK-801 (50 µM) blocked NMDAR currents in naïve cells to a similar magnitude as cells patch-loaded with MK-801 (n = 5 cells, 4 animals in each condition; U = 0.676, Mann-Whitney test). Note that CA3 pyramidal neurons were loaded for at least 3-5 minutes before recording started at +40 mV.

**Figure 3-figure supplement 1.**
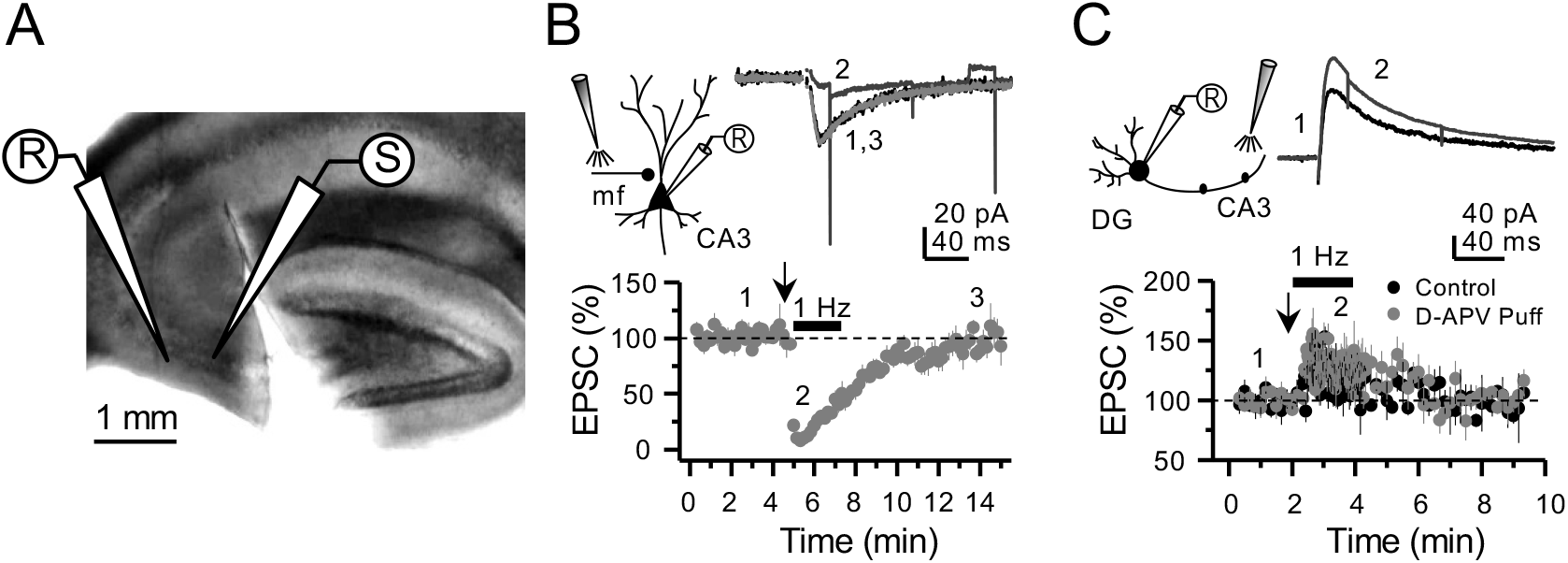
Targeting preNMDARs located in mf axons but not granule cells. (**A**) Field view of a representative hippocampal slice showing a surgical cut between DG and CA3. (**B**) Local D-APV puff application (vertical arrow, 2 puffs at 0.1 Hz) blocks NMDAR currents recorded at V_h_= -50 mV and washes out in less than 10 minutes (n = 7 cells, 5 animals, p = 5 x 10^-8^, paired *t*-test). Inset depicts the recording paradigm of the experiment *(left)*, the representative NMDAR currents *(top)* and the summary time course *(bottom)* where arrows denote the onset of D-APV (2 mM) puff application. Mfs were stimulated with a bipolar electrode (theta-glass pipette) in *stratum lucidum* in the presence of 100 µM picrotoxin and 10 µM NBQX. (**C**) D-APV puff application in CA3 did not reduce NMDAR transmission in GCs (n = 6 cells, 5 animals, control vs D-APV puff, U = 0.594, Mann Whitney test). Excitatory inputs were stimulated with a monopolar electrode placed in the medial molecular layer/inner molecular layer, in the presence of 100 µM picrotoxin and 10 µM NBQX, while GCs were clamped at V_h_=+40 mV. Data are presented as mean ± s.e.m.

**Figure 4-figure supplement 1.**
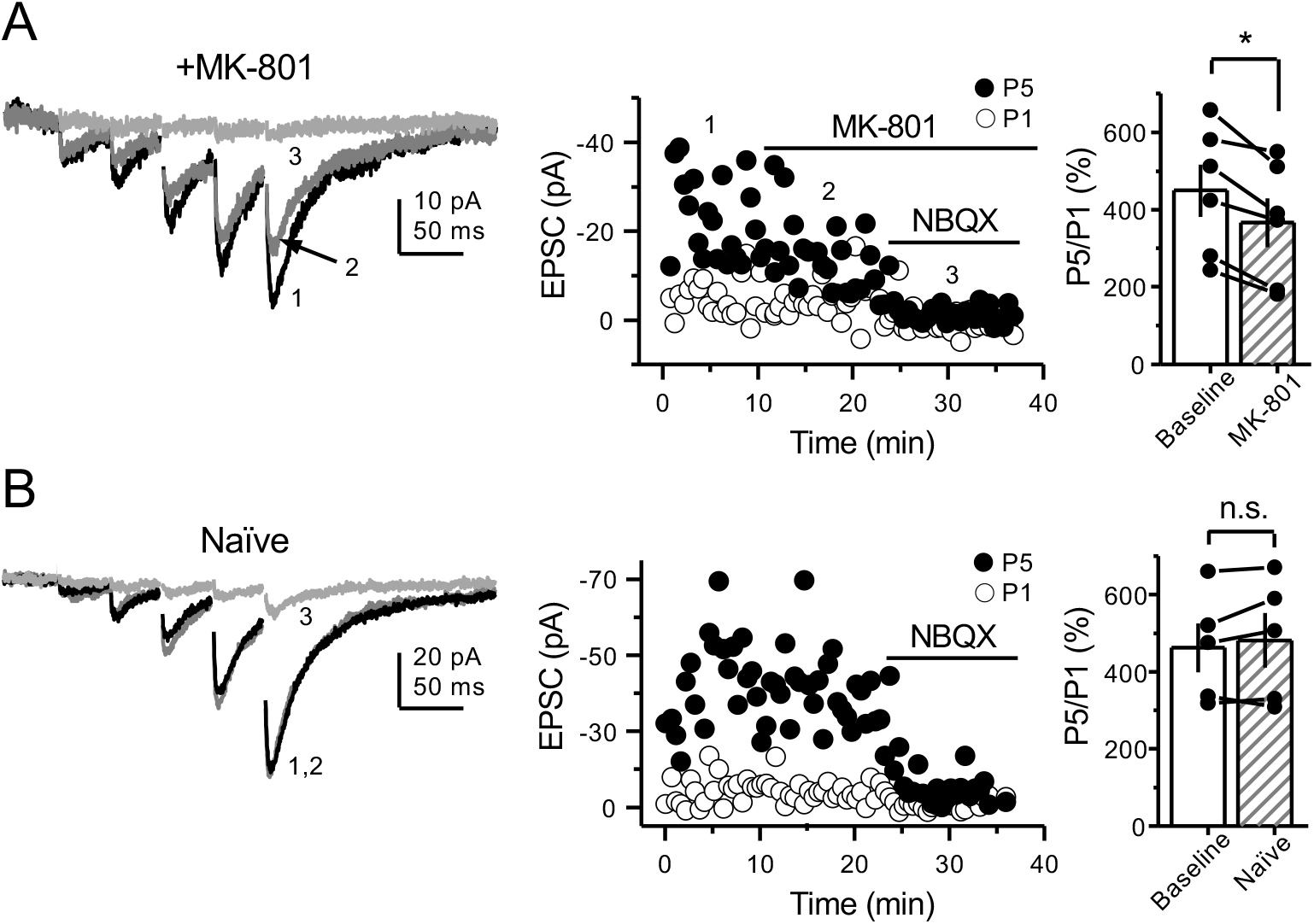
PreNMDARs contribute to burst-induced facilitation in more physiological conditions: 1.2 mM Mg^+2^, 1.2 mM Ca^+2^ and 35 °C. KAR-EPSCs were recorded from CA3 pyramidal cells loaded with 2 mM MK-801 in the presence of 15 µM LY303070 and 100 µM picrotoxin. (**A**) Bath-application of MK-801 (50 µM) significantly reduced burst-induced facilitation elicited by 5 pulses, 25 Hz (baseline 450 ± 67%, MK-801 366 ± 63%, n = 6 cells, 4 animals; baseline vs MK-801, p = 0.036, Wilcoxon Signed Ranks test). In panels A and B of this figure: representative traces (*left*), representative experiment (*middle*), and summary plot (*right*). (**B**) Burst-induced facilitation was stable in interleaved, naïve slices (baseline 462 ± 63%, naïve 481 ± 71%, n = 5 cells, 4 animals; p = 0.281, Wilcoxon Signed Ranks test). Data are presented as mean ± s.e.m. * p < 0.05.

**Figure 4-figure supplement 2.**
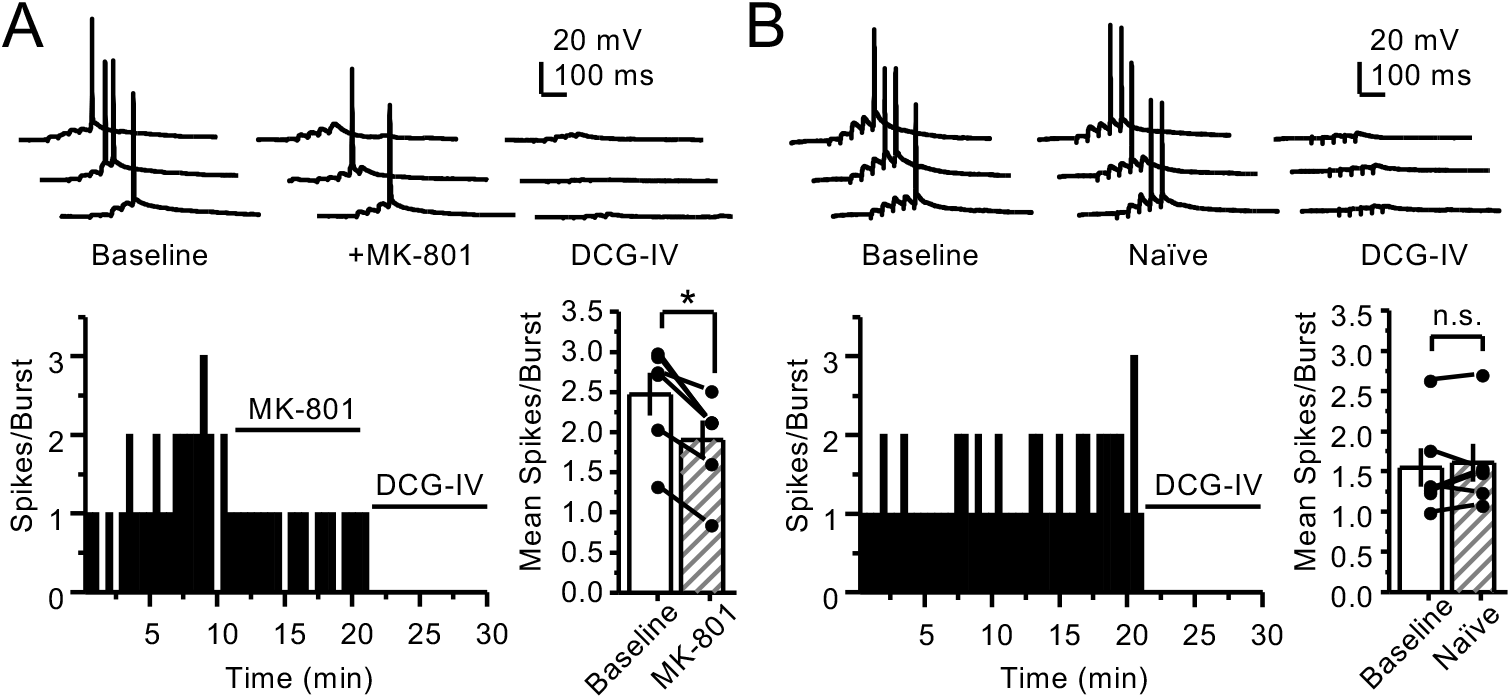
PreNMDARs contribute to action potential firing elicited by AMPAR-mediated transmission. (**A**) Bath-application of MK-801 (50 µM) reduced action potentials induced by 5 pulses at 25 Hz burst-stimulation (baseline 2.47 ± 0.27, MK-801 1.9 ± 0.24, n = 6 cells, 5 animals; p = 0.036, Wilcoxon Signed Ranks test). In panels A and B of this figure: representative traces (*top)*, representative experiment and summary plot (*bottom)*. (**B**) Stable action potential firing in interleaved naïve slices (baseline 1.55 ± 0.24, naïve 1.61 ± 0.23, n = 6 cells, 5 animals; p = 0.402, Wilcoxon Signed Ranks test). DCG-IV (1 µM) was applied at the end of all experiments in panels A-B. Data are presented as mean ± s.e.m. * p < 0.05.

**Figure 4-figure supplement 3.**
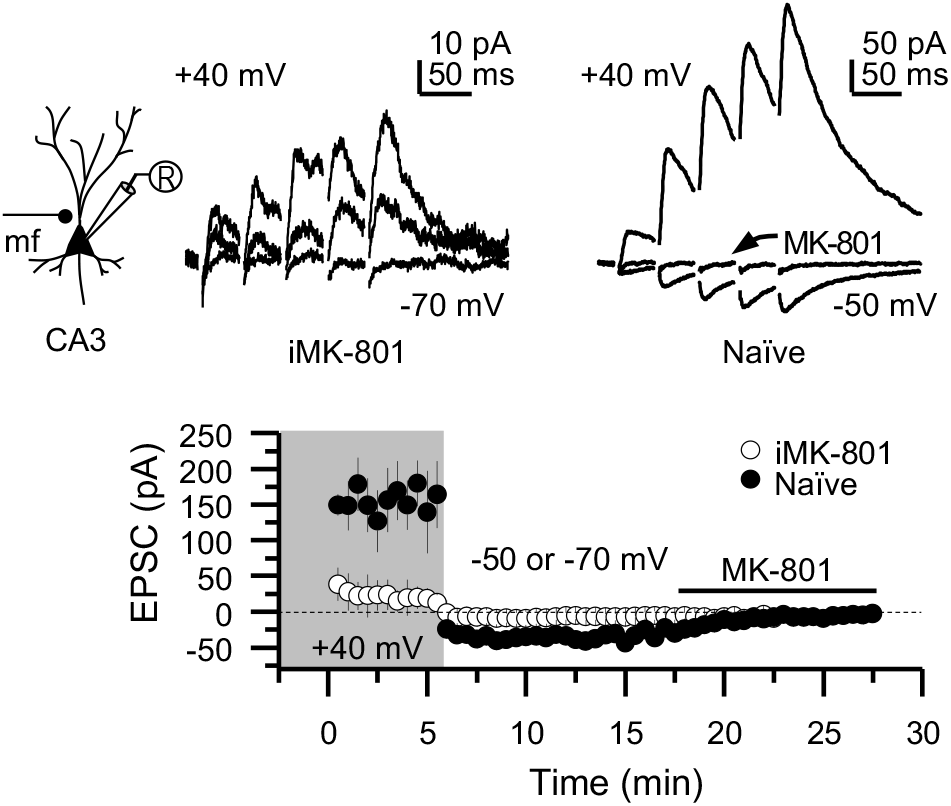
Intracellular MK-801 effectively blocked postsynaptic NMDARs elicited by burst stimulation (5 pulses at 25 Hz). Representative NMDAR-EPSCs (V_h_ = + 40 mV) from CA3 pyramidal neurons patch-loaded with 2 mM MK-801 *(left)* or naïve internal solution *(right)*. Mf inputs were stimulated with a bipolar electrode (theta-glass pipette) in *stratum lucidum* delivering in the presence of picrotoxin (100 µM) and NBQX (10 µM). NMDAR currents were recorded at V_h_= + 40 mV *(gray shaded area)* followed by a voltage jump to -70 mV in iMK- 801 conditions and -50 mV in naïve recordings. Bath-application of MK-801 (50 µM) blocked NMDAR currents of the 5^th^ pulse to a similar magnitude as iMK-801 (n = 5 cells, 4 animals per condition; U = 0.21, Mann-Whitney test). Data are presented as mean ± s.e.m.

**Figure 5-figure supplement 1.**
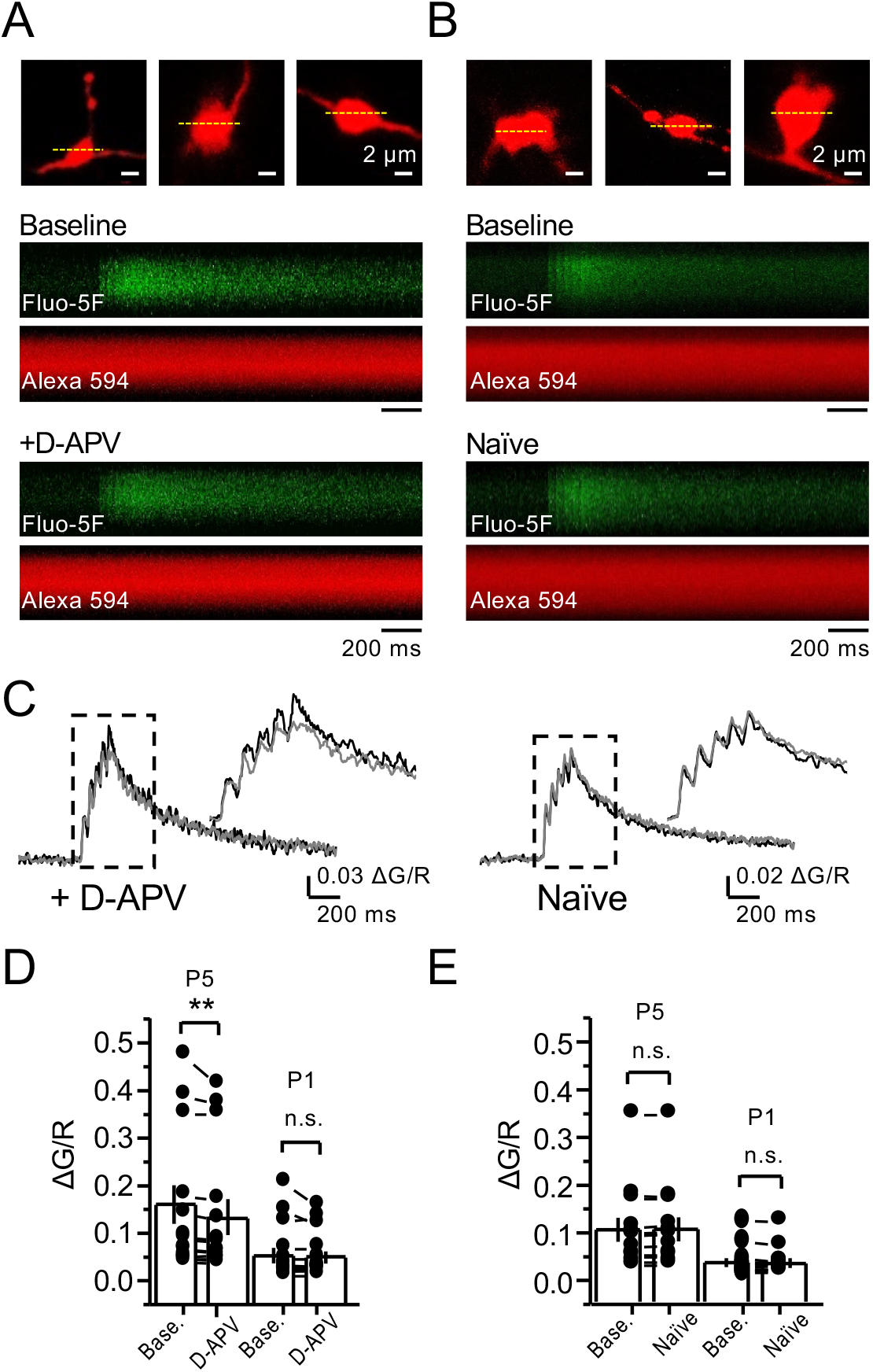
NMDAR antagonism reveals a reduction in presynaptic Ca^+2^ rise in the presence of 1.3 mM Mg^+2^ and 2.5 mM Ca^+2^. (**A, B**) Granule cells were patch-loaded with Fluo-5F (200 µM) and Alexa 594 (35 µM). Line scan analysis of mf giant bouton calcium transients (CaTs) in response to action potential (AP) stimulation (5 APs, 25 Hz). (**C**) Line scan signals following D-APV application or naïve conditions. (**D**) D-APV (100 µM) significantly reduced the 5^th^ peak (P5) of CaTs (baseline 0.155 ± 0.04, D-APV 0.138 ± 0.03, n = 13 boutons, 10 animals; baseline vs D-APV, p = 0.00642, Wilcoxon Signed Ranks test). (**E**) In naïve conditions P5 of CaTs is stable (baseline 0.104 ± 0.026, naïve 0.105 ± 0.026, n = 12 boutons, 10 animals; baseline vs naïve, p = 0.255, Wilcoxon Signed Ranks test). The first peak (P1) of CaTs is not affected by D-APV (baseline 0.05 ± 0.017; D-APV 0.047 ± 0.014; baseline vs D- APV, p = 0.485, Wilcoxon Signed Ranks test) and is stable in naïve conditions (baseline 0.033 ± 0.009; naïve 0.032 ± 0.009, baseline vs naïve, p = 0.196, Wilcoxon Signed Ranks test). Data are presented as mean ± s.e.m. ** p < 0.01.

**Figure 6-figure supplement 1.**
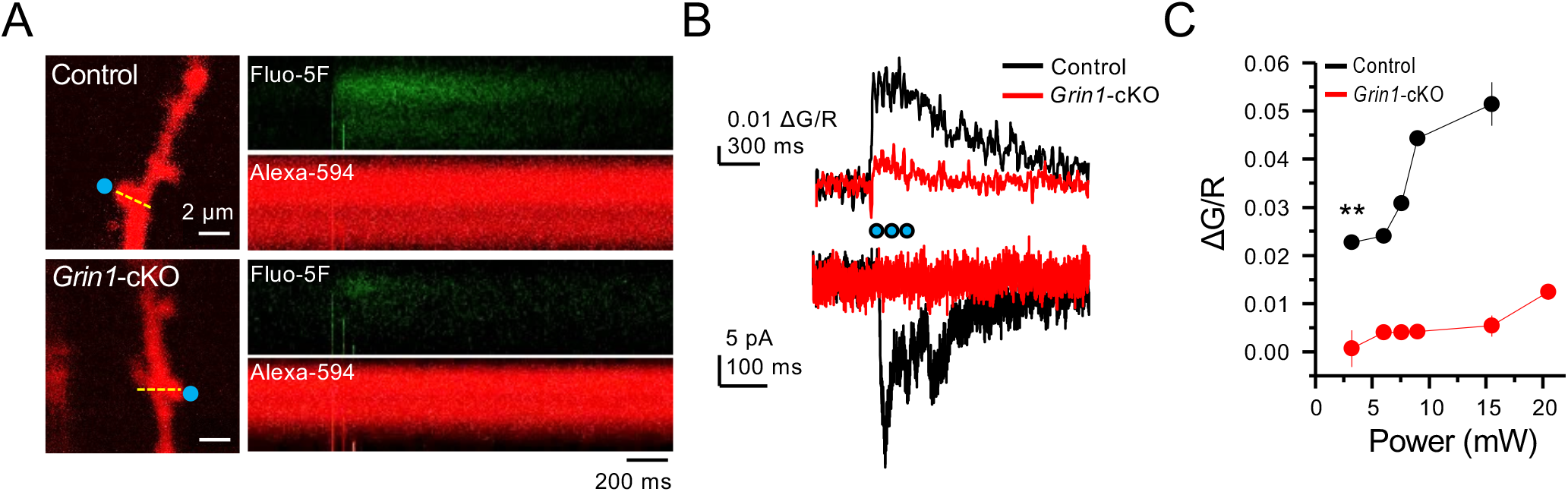
*Grin1*-cKO exhibit reduced CaTs at varying uncaging laser power intensities. (**A**) Representative images of CaTs from control *(top)* and G*rin1*-cKO animals *(bottom)* after MNI-glutamate uncaging (2 mM, 3 pulses at 25 Hz) on GC dendritic spines. Dotted line (yellow) indicates line scan, and blue dots indicate 2PU spots. (**B**) Quantified ΔG/R signals *(top)* and uncaging induced NMDAR-EPSCs *(bottom)* from control and *Grin1*-cKO animals. Blue dots indicate when 2PU pulses were delivered. (**C**) Control animals display robust ΔG/R signals as compared to *Grin1*-cKO animals at varying laser power intensities (6 spines, 6 animals per group, U = 0.00507 per power intensity, Mann-Whitney test). Data are presented as mean ± s.e.m. ** U < 0.01.

**Figure 6-figure supplement 2.**
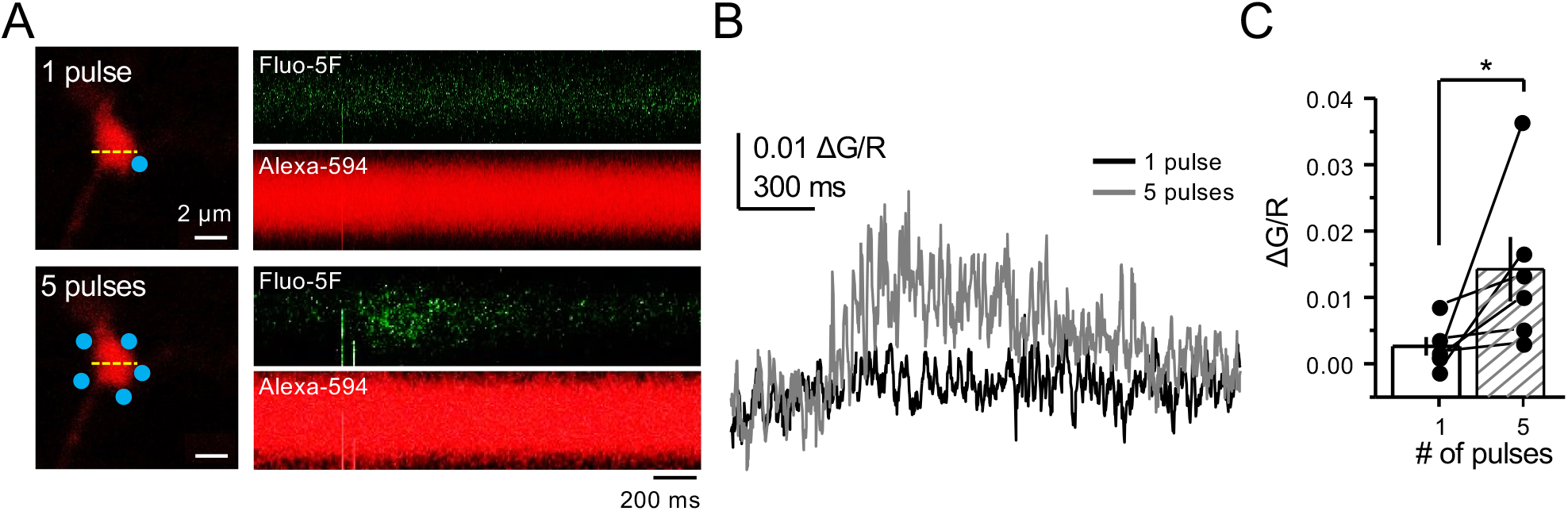
Bouton CaTs can be detected after repetitive uncaging of MNI- glutamate. (**A**) Representative images of CaTs from single-trial: 1 pulse *(top)* and 5 pulses, 25 Hz *(bottom)* of MNI-glutamate uncaging (2 mM). Dotted line (yellow) indicates line scan, and blue dots indicate 2PU spots. (**B**) Quantified ΔG/R signals from 1 pulse *(black)* and 5 pulses *(dark gray)* from all trials. (**C**) Repetitive pulses result in larger ΔG/R signals as compared to single pulses (n = 6 boutons, 6 animals, p = 0.03603, Wilcoxon-Signed Ranks test). Data are presented as mean ± s.e.m. * p < 0.05.

**Figure 6-figure supplement 3.**
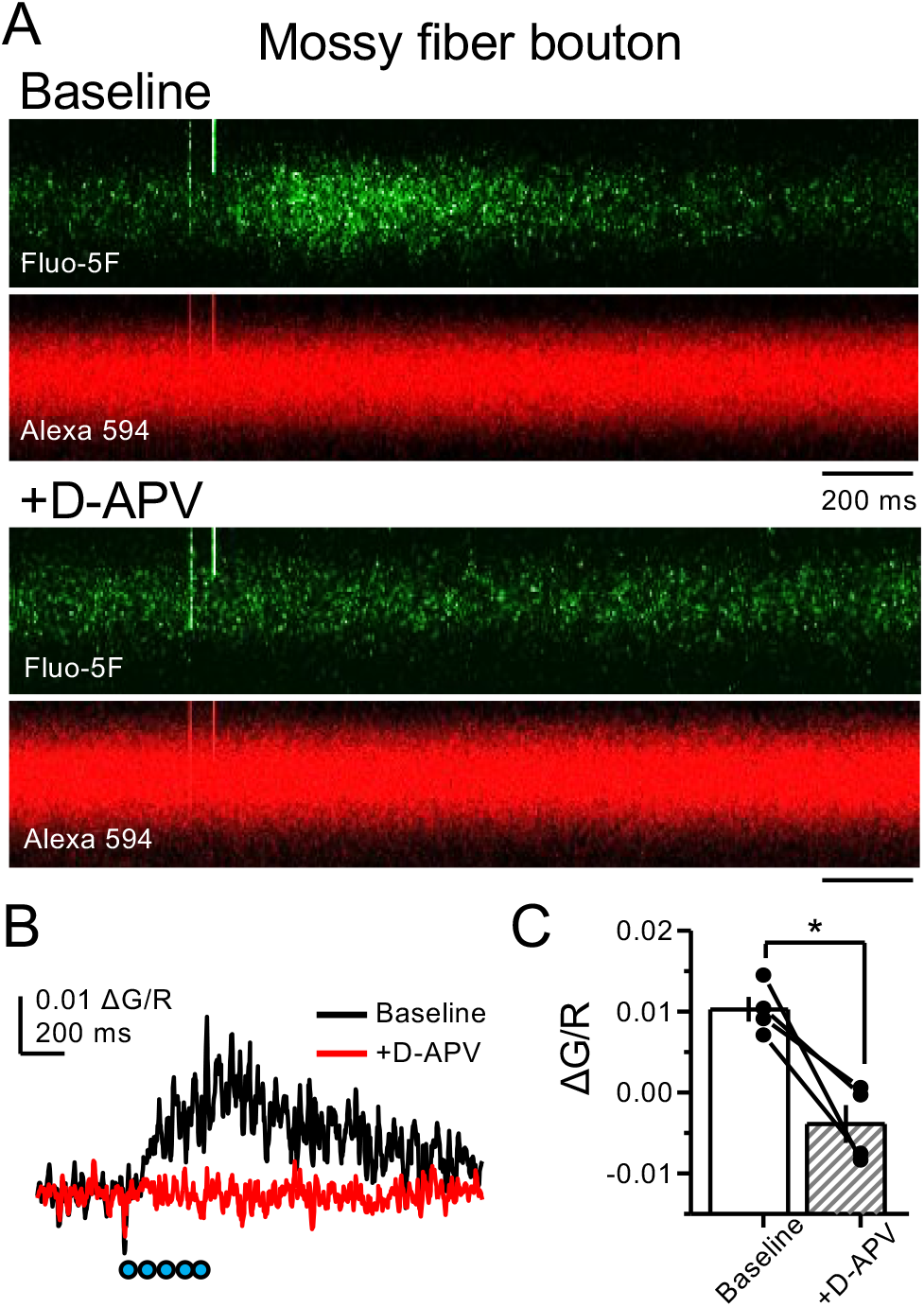
NMDAR antagonism with D-APV blocks CaTs elicited by glutamate 2PU. (**A**) Representative image of baseline glutamate uncaging driven CaTs in mf boutons *(top)*. D-APV application (100 µM) blocks CaTs *(bottom)*. (**B**) Quantified ΔG/R signals before and after D-APV application. (**C**) Summary data of D-APV block on glutamate uncaging elicited CaTs (baseline 0.0103 ± 0.0016, D-APV -0.004 ± 0.0024, n = 4 boutons, 3 animals; U = 0.0304, Mann-Whitney test, baseline vs D-APV). Data are presented as mean ± s.e.m. * U < 0.05.

**Figure 7-figure supplement 1.**
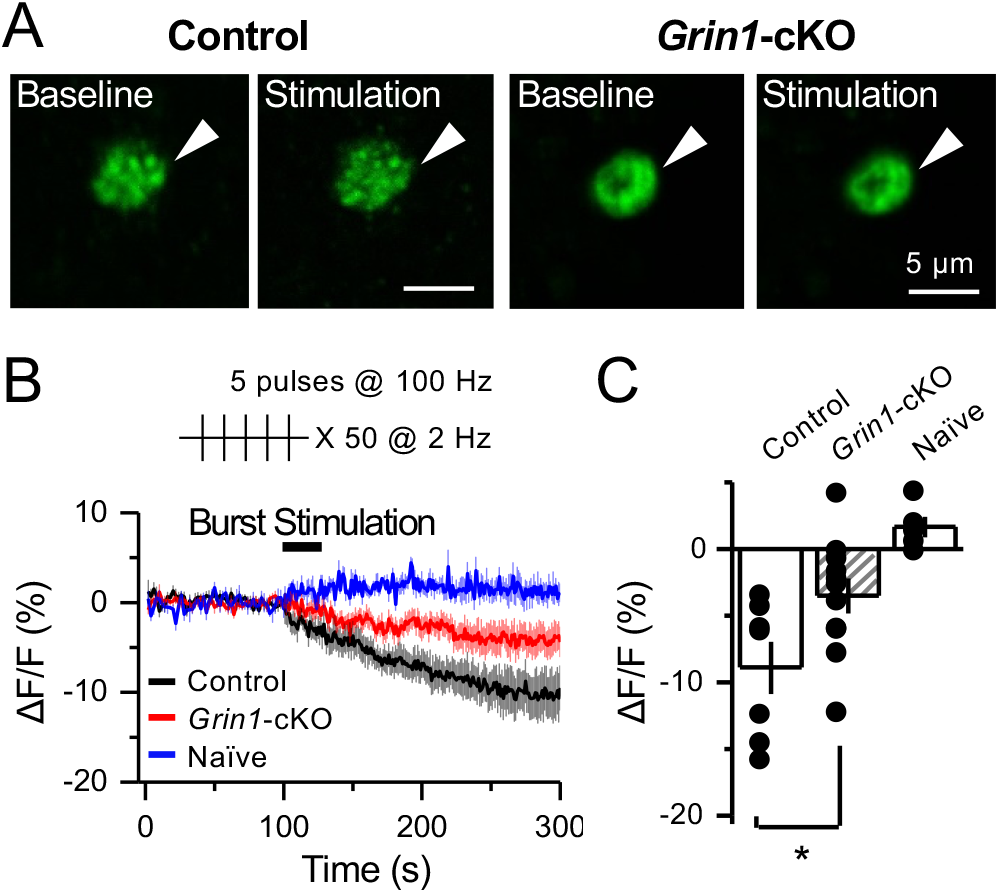
preNMDARs contribute significantly to BDNF release following a more physiological pattern of burst-stimulation. (**A**) Representative images of BDNF-pHluorin signal intensity at baseline and after burst stimulation of mfs (5 pulses, 100 Hz, x 50, every 0.5 s). Control images *(left)*, *Grin1*-cKO images *(right*), arrowhead indicates region of interest. (**B**) Time course of BDNF-pHluorin signal intensity measured as ΔF/F (%): control *(black)*, *Grin1*- cKO *(red),* Naïve *(blue)*. (**C**) Quantification of BDNF-pHluorin signal in (B) during the last 100 seconds reveals larger BDNF release in control animals as compared to *Grin1*-cKO (control - 8.9% ± 2%, n = 7 slices, 5 animals; *Grin1*-cKO -3.5 ± 1%, n = 11 slices, 5 animals; *Grin1*-cKO vs. control, p = 0.0305, unpaired *t*-test). Data are presented as mean ± s.e.m. * p < 0.05.

**Figure 8-figure supplement 1.**
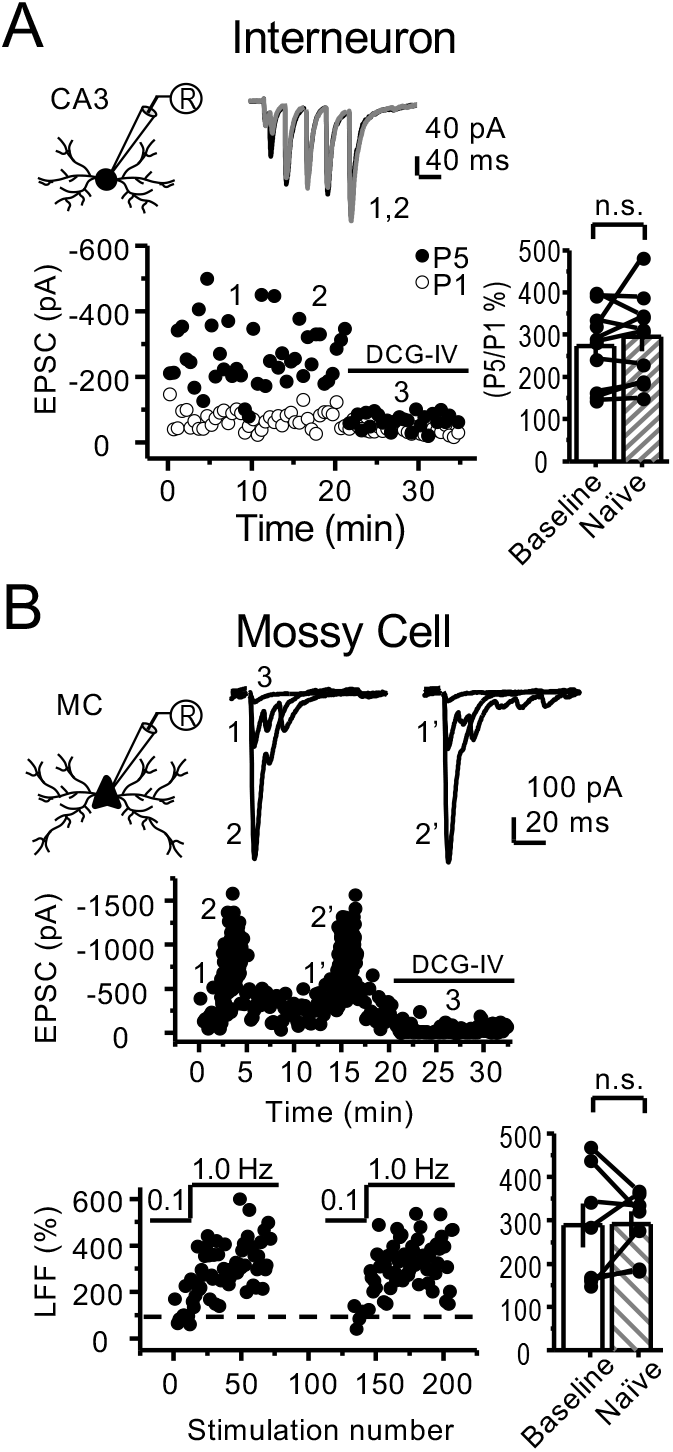
Stability experiments for mf-Interneuron and mf-mossy cell short-term plasticity. (**A**) Stable CA3 interneuron burst-induced facilitation of mf-CA3 transmission (baseline 273 ± 30%, naïve 294 ± 33%, n = 10 cells, 6 animals; p = 0.298, paired *t*- test, baseline vs naïve). (**B**) Stable low-frequency facilitation (LFF) of AMPAR-EPSCs in hilar mossy cells (baseline 288 ± 51%, naïve 291 ± 29%, n = 7 cells, 6 animals; p = 0.937, paired *t*- test, baseline vs naïve). DCG-IV (1 µM) was applied at the end of all recordings to confirm mf- CA3 transmission. Data are presented as mean ± s.e.m.

## Acknowledgements

We thank all the Castillo lab members for invaluable discussions. We also thank Dr. Hyungju Park for his generous gift of the BDNF-phluorin DNA construct, Dr. Michael Higley for sharing *Grin1* floxed mice, and Dr. Pascal Kaeser for his generous gift of the Cre-dependent ChIEF DNA construct.

## Key Resources Table

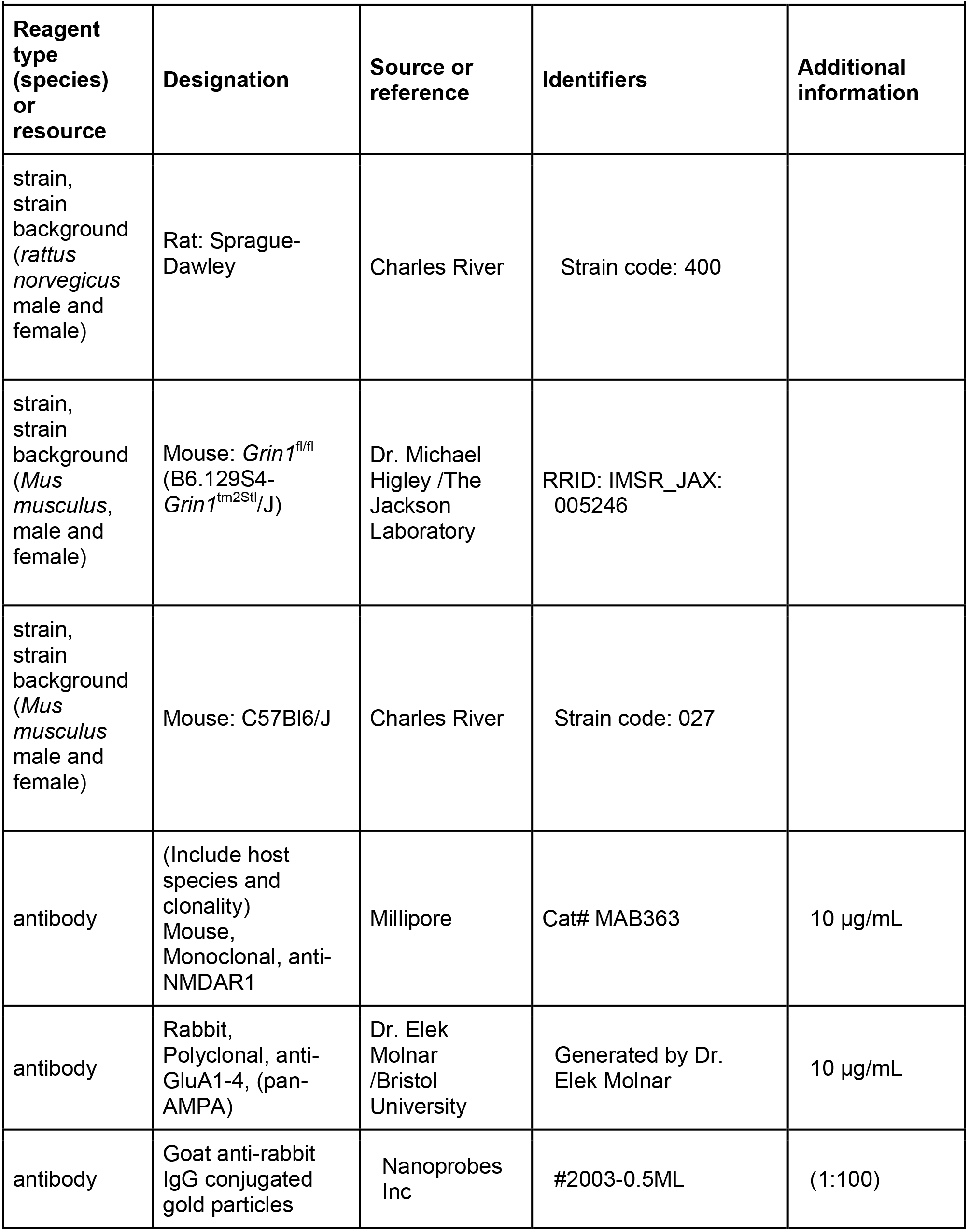
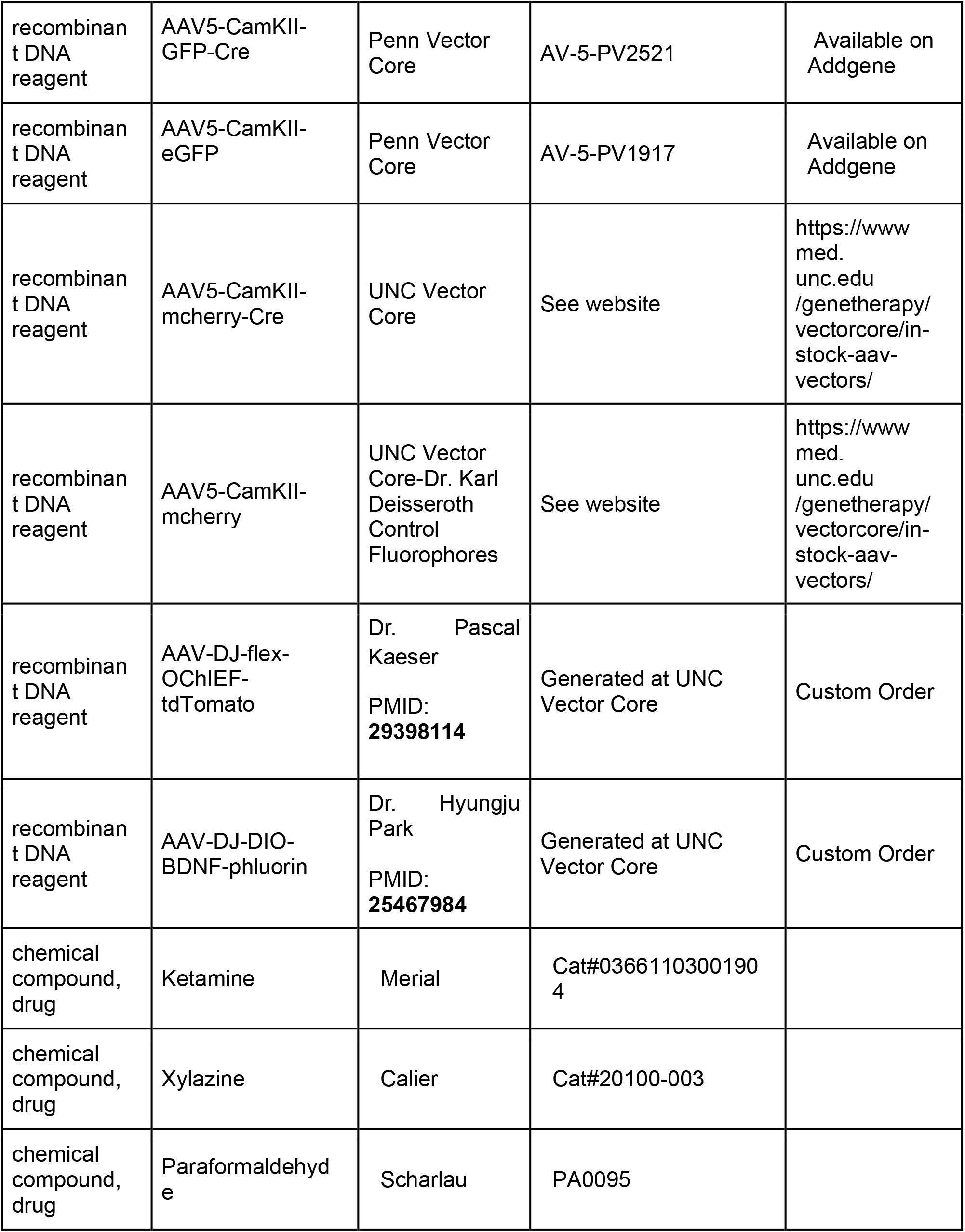
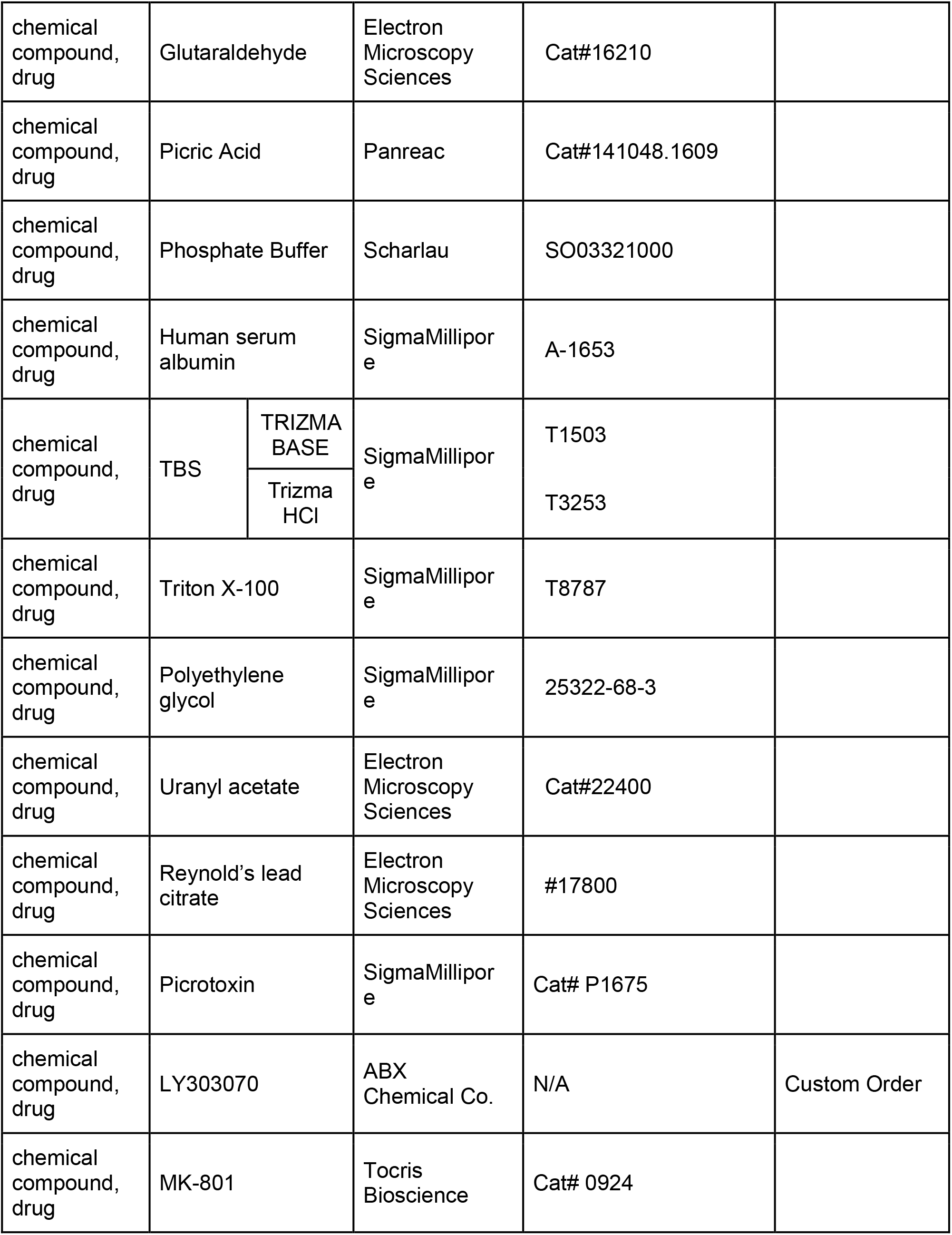
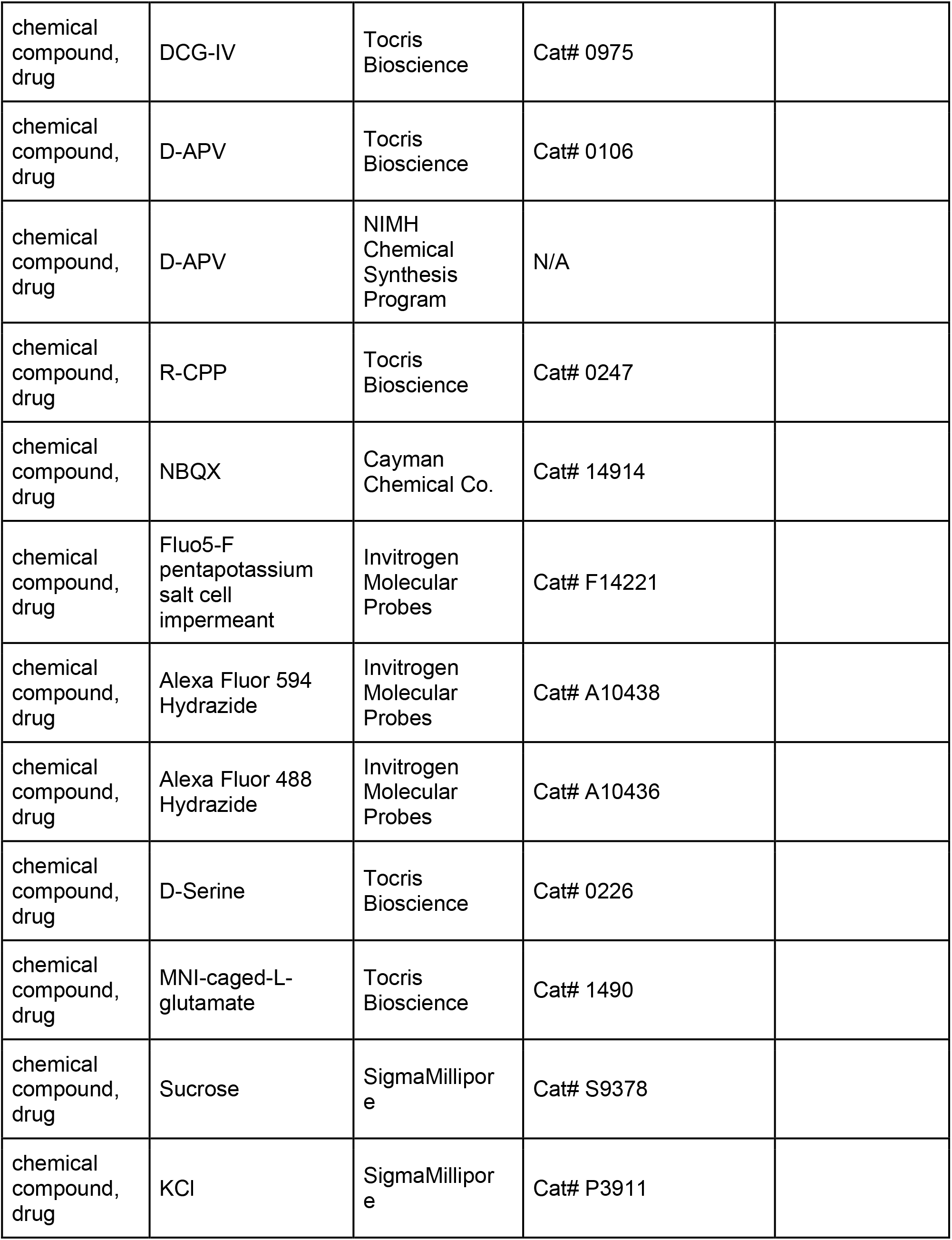
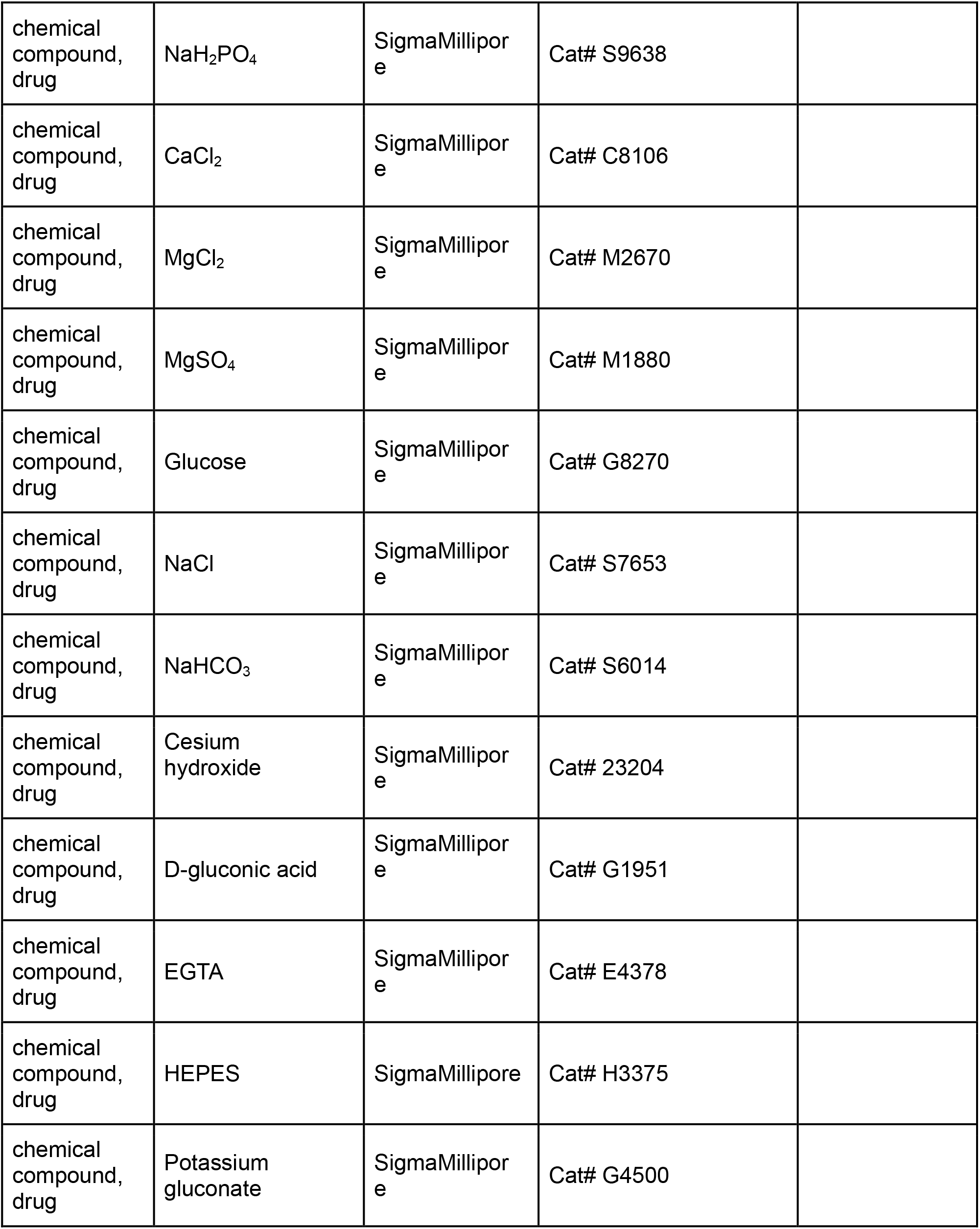
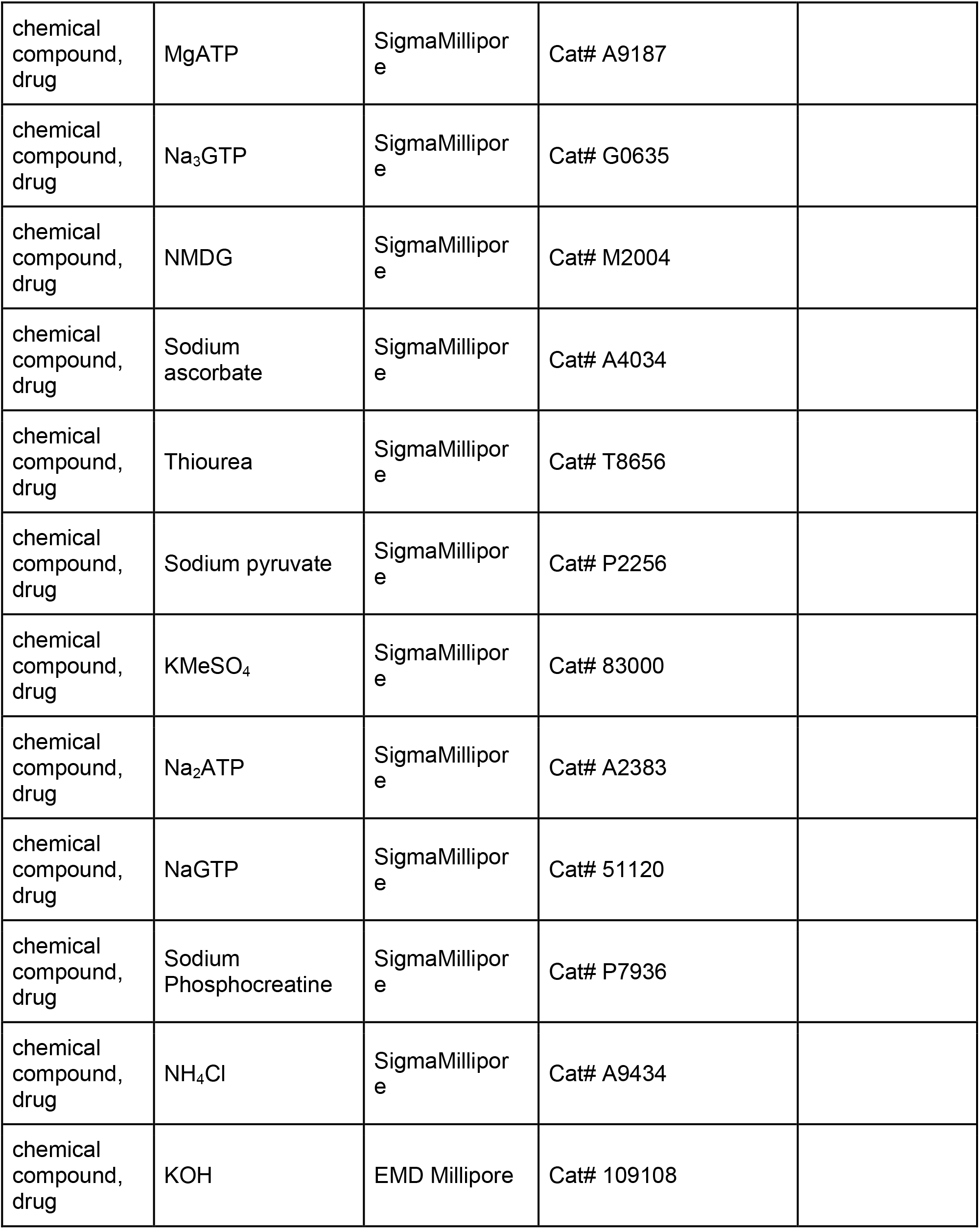
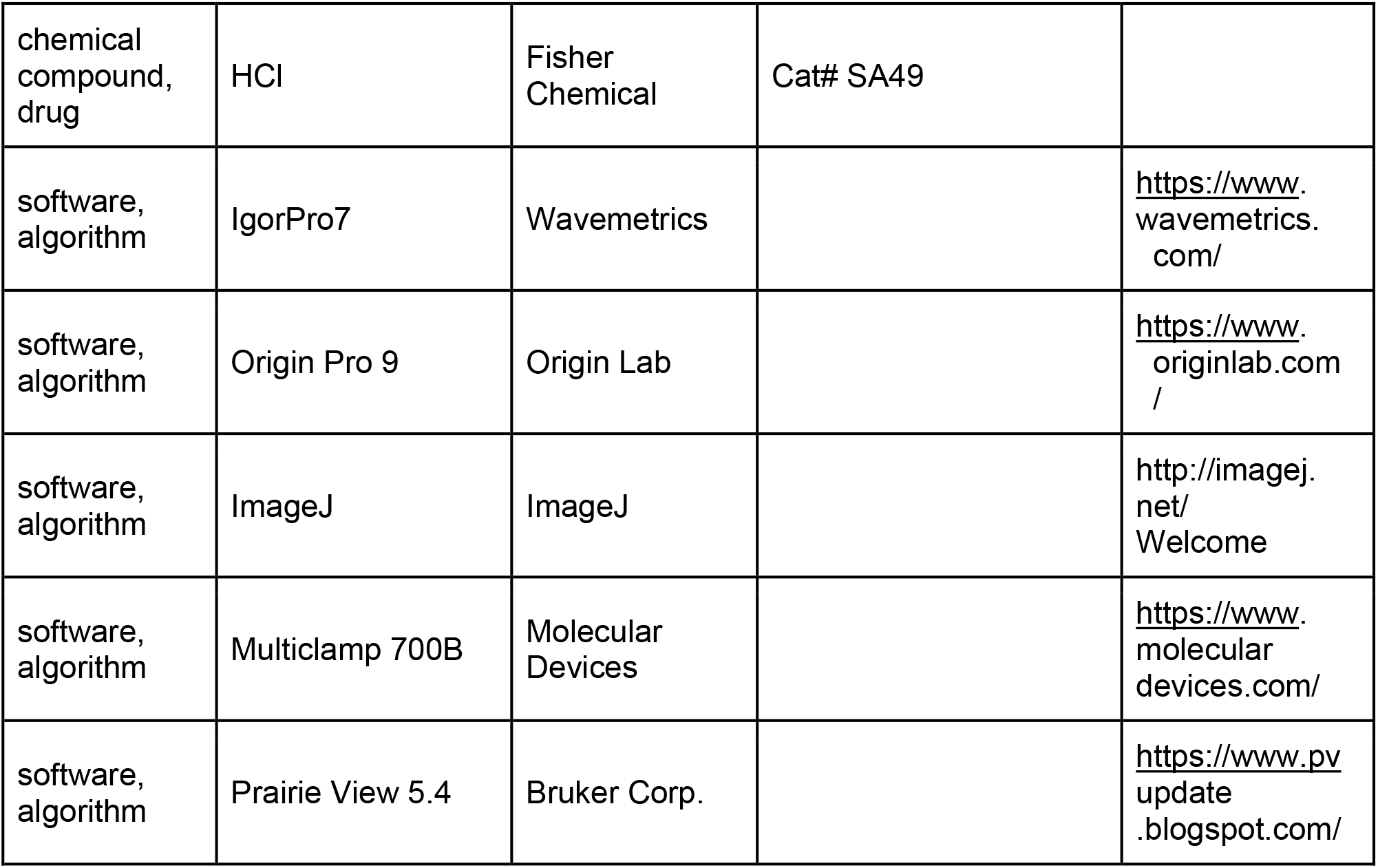

